# Bite force transmission and mandible shape in grasshoppers, crickets, and allies is largely dependent on phylogeny, not diet

**DOI:** 10.1101/2023.03.28.534586

**Authors:** Carina Edel, Peter T. Rühr, Melina Frenzel, Thomas van de Kamp, Tomáš Faragó, Jörg U. Hammel, Fabian Wilde, Alexander Blanke

**Author notes:** Corresponding author / Carina Edel.

## Abstract

Although organ systems evolve in response to many intrinsic and extrinsic factors, frequently one factor has a dominating influence. For example, mouthpart shape and mechanics are thought to correlate strongly with aspects of the diet. Within insects, this paradigm of a shape-diet connection is advocated for decades but the relationship has so far never been quantified and is mostly based on qualitative observations. Orthoptera (grasshoppers, crickets, and allies) are a prominent case, for which mandible shape and dietary preference are thought to correlate strongly and even lead to predictions of feeding preferences. Here, we analysed mandible shape, biting efficiency, and their potential correlation with dietary categories in a phylogenetic framework for a broad sampling of several hundred extant Orthoptera covering nearly all families. The mandibular mechanical advantage was used as a descriptor of gnathal edge shape and bite force transmission efficiency. We aimed to understand how mandible shape is linked to biting efficiency and diet, and how these traits are influenced by phylogeny and allometry. The investigation reveals that feeding ecology is not the unequivocal predictor of mandible shape that it was assumed to be. There is a strong phylogenetic signal suggesting that phylogenetic history does have a much more prevalent influence on gnathal edge shape and distal mechanical advantage, than, e.g., feeding guilds or the efficiency of the force transmission to the food. Being ancestrally phytophagous, Orthoptera evolved in an environment with abundant food sources so that selective pressures leading to more specialized mouthpart shapes and force transmission efficiencies were low.

## Introduction

A conjecture in biology is that form and function vary together in a correlated pattern. This lead to the assumption that shape might even predict function and, consequently, aspects of the ecological niche (Feilich and López-Fernández, 2019). For example, in 1862 Darwin predicted the existence of a hawkmoth, Xanthopan morgani, with an extremely elongated tongue, because he discovered an orchid with an extremely elongated nectar tube (Burkhardt et al., 1985). Years later, in 1903, Darwins’ prediction was proven (Rothschild Jordan 1903). In this context, mouthpart phenotypes are examples of tight adaptations to food sources with remarkable patterns of convergence. Jaw shape and dentation show convergence in mammals in distantly related orders such as the aye-aye (Daubentonia madagascariensis) and the squirrels (Scirius) (Berthaume et al., 2019; Evans et al., 2007; Grossnickle, 2020; Morales-García et al., 2021; Morris et al., 2018). A correlation between diet and jaw shape was also found in non-mammal vertebrate orders such as fishes (Carroll et al., 2004; Wainwright and Richard, 1995), lizards (Metzger and Herrel, 2005), and birds (Olsen, 2017).

While feeding performance is an integral part of survival (Stephens and Krebs, 2019) and an increase in efficiency through shape adaptation may be advantageous, it may be overshadowed by a complex interplay of behavioural, evolutionary and physiological factors and, in addition, mechanical constraints. A lack of an ecomorphological correlation between diet and function may be the result.

Mouthpart shape disparity is immense in chewing-biting insects and its relationship to diet is considered to be very tight. This was established by qualitative studies which investigated Orthoptera as well as some other polyneopterans (Aguirre-Segura et al., 1987; Bennack, 1981; ElEla et al., 2010; Gangwere, 1965; Isely, 1944; Kang et al., 1999; Kaufmann, 1965; Patterson, 1984; Samways et al., 1997; Smith and Capinera, 2005). With ∼30.000 species Orthoptera are the most diverse non-holometabolan biting-chewing insect group. They are mostly phytophagous insects and can, have immensely damaging effects on plants including crops. Diet preferences can range from very specialised in form of phytophage monophagy (e.g. Bootettix sp. Otte and Joern, 1976) and obligate carnivory (e.g. Saga pedo), to omnivory (Gangwere, 1961; Ingrisch and Köhler, 1998). Feeding mode is equally diverse, including sedentary grazing (most Acrididae), scavenging (e.g. Gryllidae) and active predation (e.g. Saga pedo) (Kaltenbach, 1990; Lupu, 2007). More uncommon food sources are found within the subfamily Zaprochilinae, which feed on pollen and are a rare example of pollinating Orthoptera (Tan et al., 2017). With all those different organic materials, nutritional compositions such as protein-to-carbohydrate ratios, vary between plant and animal food sources and specialization to one limits access to a nutrient-diverse diet. Herbivores e.g. have access to abundant food resources but no easy protein source especially in grass-feeders (Hochuli, 1996; Le Gall and Behmer, 2014). Food toughness also varies ranging from rather softer animal sources, like worms and larvae, to tougher plant matters like grasses (Clissold, 2007; Clissold et al., 2009; Schoonhoven et al., 2005). For the majority of herbivorous animals fractioning plant material with their teeth is the key factor affecting nutrient uptake (Sanson, 2006) and its efficiency can be associated with mandible morphology (Bennack, 1981).

Mandible shape in orthopterans was categorized into dietary preference types (Gangwere, 1966; Isely, 1944) such as graminivorous, forbivorous, carnivorous, detritivorous and omnivorous, with some in-between forms mentioned. Based on such qualitative descriptions authors inferred diet preference based on mandible shape (ElEla et al., 2010; Gangwere and Spiller, 1995; Smith and Capinera, 2005). The different mandible types show variation in the geometry of the gnathal edge, the molar region and in width to length ratio. Similarly to mammals, such geometric adaptations are thought to increase nutrient uptake because populations adapt to the hardness and material composition of the target food (Bernays et al., 1991; Gangwere, 1965; Patterson, 1984, 1983; Püffel et al., 2021; Weihmann et al., 2015). Shorter stouter mandibles with a clearly defined and ridged molar area were qualitatively linked to tougher plant matters like grasses (Gangwere et al., 1998; Kaufmann, 1965). On the other end of this extreme are carnivorous mandibles, which were described as elongated with a hook-like shape and a flat, non-structured molar area. Elongation is linked with increased biting speed and is thought to be adapted to prey capturing (Corbin et al., 2015; Stayton, 2006; Westneat, 2004). Between those two extremes exists an immense variation of in-between forms with varying degrees of the tooth and molar definition (Gangwere, 1965; Isely, 1944). Assigning those intermediate shapes to different dietary categories has been attempted (Gangwere, 1965; Ingrisch and Köhler, 1998; Isely, 1944; Kaufmann, 1965; Uvarov, 1966), but a clear and common definition is missing. Here, we use the mechanical advantage as a biomechanical performance metric and shape descriptor for the orthopteran mandible to determine if diet and gnathal edge shape follow the presumed one-to-one mapping in a phylogenetic framework.

## Material and Methods

### Taxon sampling

We studied 337 species of Orthoptera from 316 different genera, covering all extant families, except Pyrgacrididae, Cylindrachetidae, and Cooloolidae (S 11). Specimens from the orders Dermaptera, Blattodea, Plecoptera, Zoraptera, Grylloblattodea and Phasmatodea were used as outgroups. All specimens were dried museum specimens from the National History Museum (NHMUK, London, UK), Zoologisches Forschungsmuseum Koenig (ZMFK, Bonn, Germany), Naturhistorisches Museum (NHMV, Vienna), Museum fuer Naturkunde (MfN, Berlin, Germany), and Zoologische Staatssammlung (ZSM, Munich, Germany).

### Ecological data sampling

Eight different diet guilds were defined, based on the most common literature mentions (Tab. 1). Using the software ‘Publish or Perish’ (v: 7) (Harzing, 2007) a Google Scholar search was conducted for each species. Each of the ∼2000 publications were searched for diet information (using the keywords ‘diet’, ‘food’, and ‘feeding’) and a feeding category was assigned for each species. If no diet information for a species could be found, similar searches were conducted using synonyms, genus, or subfamily status. All diet information from the Orthoptera Species File (OSF) (“Orthoptera Species File v. 5.0/5.0.,” 2021) (retrieval date 13. April 2021) was extracted and cross-referenced with the literature data. If a mismatch occurred the literature data got preference. If no literature data was found the information was supplemented with OSF data.

**Tab. 1:**
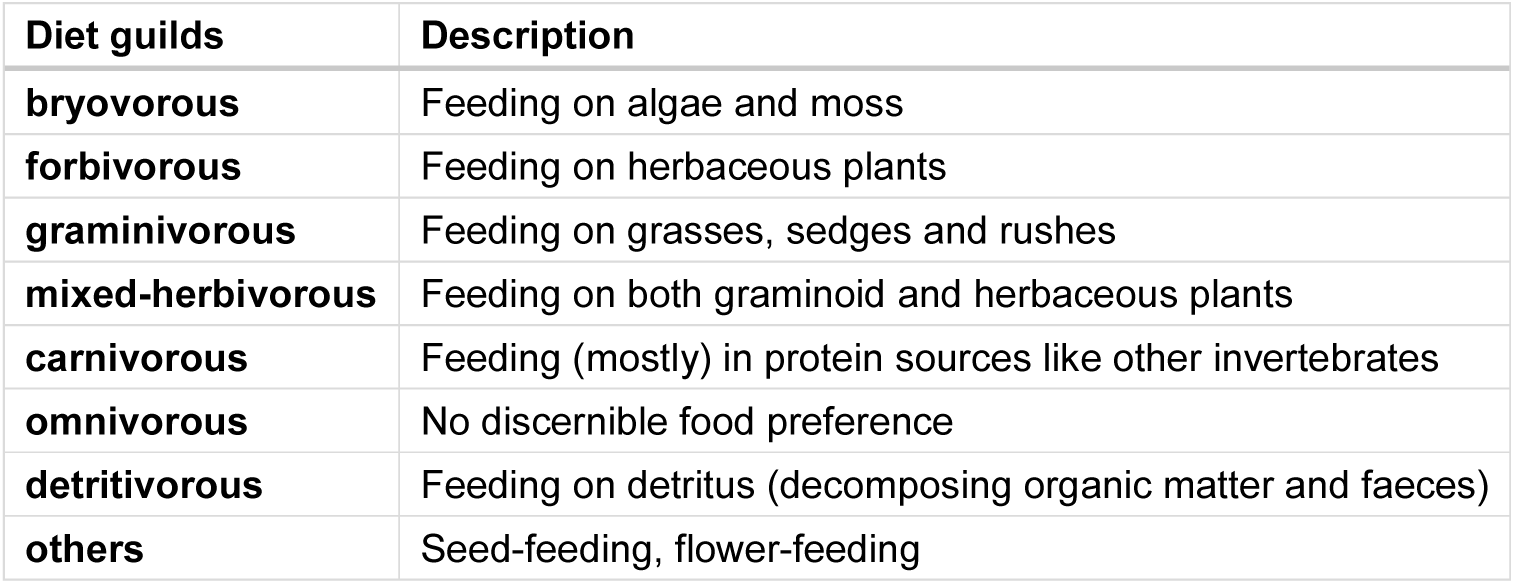
Definition of diet guilds based on most common literature mentions

### µCT-Scanning and 3D reconstruction

The heads of all specimens were scanned with synchrotron radiation micro-computed tomography (SR-µCT) at different imaging facilities. 98 scans were done at KIT Light Source (Karlsruhe Institute of Technology (KIT), Karlsruhe, Germany), 44 at TOMCAT (Stampanoni et al., 2006) (Swiss Light Source (SLS), Paul-Scherrer-Institute (PSI), Villigen, Switzerland), and 21 at the Center for Biohybrid Medical Systems (CBMS) (Aachen, Germany). A further 190 specimens were scanned at DESY (Deutsches Elektronen Synchrotron, Hamburg, Germany) of which 106 were processed at the IBL-P05 imaging beamline (Khokhriakov et al., 2017; Moosmann et al., 2014; Wilde et al., 2016) (operated by the Helmholtz-Zentrum Hereon at PETRA III) and 74 larger specimens at the Phoenix Nanotom, General Electric, Boston, MA) housed at DESY. Each scan was downsampled to ∼300 MB with Fiji (Schindelin et al., 2012) using a stack-cropping macro script of Rühr et al., 2021, which also generates ‘HDR5-Analyse’ files with a corresponding .hdr file for import into ITK-Snap (v. 3.8) (Yushkevich et al., 2006). 3D reconstruction was done in ITK-Snap with a pre-segmentation by hand and completed using a semi-automatic segmentation algorithm. Miss-assigned voxels were afterwards corrected by hand and a smooth .stl surface was exported for import into Blender (v. 3.8) (Hess, 2010).

### Mechanical advantage and the bite efficiency profile

Orthoptera have a dicondylic mandible which articulates with the head capsule via two ball-and-socket joints. The mandibles, therefore, rotate around only one axis going through the centres of those two joints creating a virtual hinge joint. The mandibles are mainly moved by two muscles, a closer muscle which occupies most of the head volume and a much smaller opener muscle (Chapman, 1995; Snodgrass, 1935). The mandibles are slightly asymmetrical with the left mandible overlapping the right mandible and both mandible’s biting areas fitting together in a key lock principle (Chapman, 1995; Snodgrass, 1935). Because of this, only the left mandible is used in all analyses. The biting area of the mandibles differentiates into a distal incisor lobe (pars incicivus) and a proximal molar lobe (pars molaris) (Chapman, 1995; Richter et al., 2002). Together the lobes form the gnathal edge of the mandible (Edgecombe et al., 2003).

The effectivity of the force transmission from the muscles via the mandibles to the biting area can be described with mechanical advantage (MA) which is the ratio of in-lever to out-lever length (Clissold, 2007; Westneat, 2004). For insect mandibles, the in-lever is the perpendicular distance between the fulcrum and muscle insertion point, whereas the out-lever is the distance between the fulcrum and the biting point (Fig. 1).

**Fig. 1:**
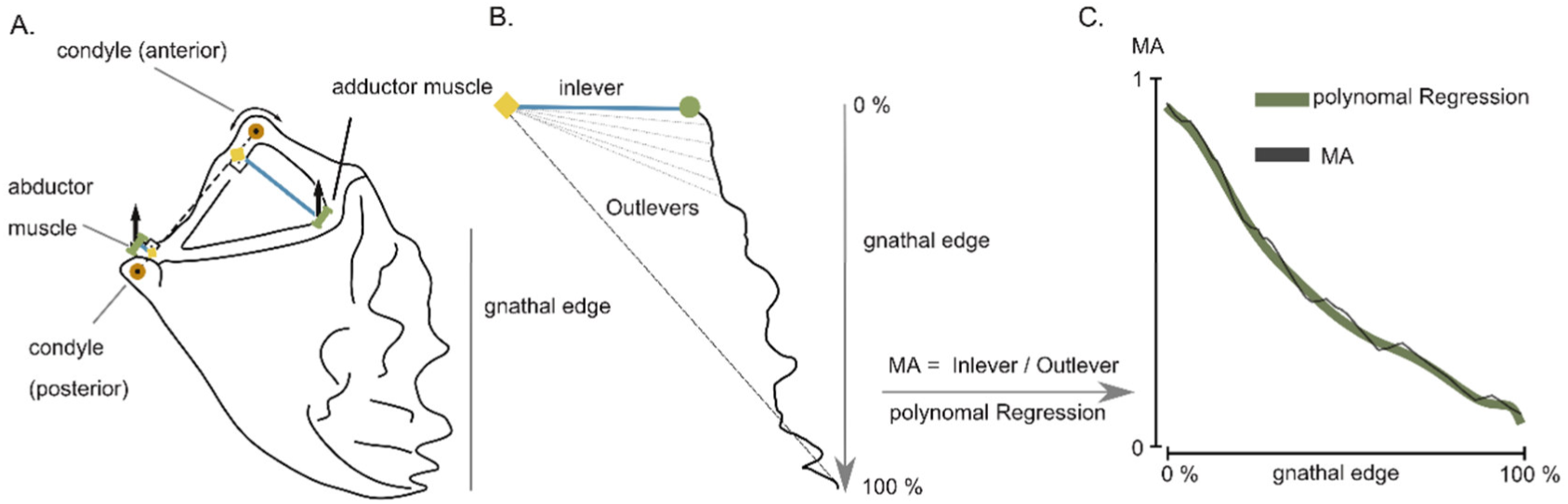
Methodological overview for mechanical advantage measurements and their translation into bite efficiency profiles. *A. A schematic of a typical mandible with the rotation axis point (yellow) between the joints (orange) and the in-lever (blue)*. *B. Gnathal edge measurements and calculation of mechanical advantage for a rotated mandible so that the rotation axis is perpendicular to the plane of projection. In-lever with the point of rotation (yellow) and closer muscle insertion point (green)*. *C. Example of the mechanical advantage (MA) progression along the gnathal edge (black) and the resulting polynomial curve (green)*.

3D reconstructed mandibles were used for MA measurements with Blender and R. After import in Blender the mandibles were rotated so that the rotation axis of all mandibles was aligned on the same plane and the in-lever for the adductor and abductor muscle start and endpoint 3D coordinates exported. The mandibles were flattened along the sagittal midline and a line with 400 to 800 vertices wrapped around the gnathal edge (Fig. 1). 3D coordinates were exported from Blender into R (v. 4.2.1) (R Core Team, 2022) and the in-lever and out-lever lengths were calculated for both mandible muscles. Because clearly defining the start of the molar area was often impossible the biting area was measured from the adductor muscle insertion point to the most distal incisivi and normalized with a scale from 0% to 100%. The mechanical advantage was then calculated for each point along the gnathal edge which then forms the bite efficiency profile (BEP). A polynomial function was fitted through the MA progression along the tooth row for each specimen. The best fit for the polynomial function was determined based on the Akaike (AIC) and Bayes (BIC) information criteria (Bozdogan, 1987). Polynomial models from 1 to 20th degree were tested with the function best_fit() in forceR (Rühr and Blanke, 2022) to determine the AIC value which changes less than 5% from one value to the next. Further statistical tests were then conducted with the coefficients of the polynomial curve with the best fit according to the AIC.

### Phylogenetic comparative methods

The phylogenetic signal was determined using the most recent and comprehensive time-calibrated Orthoptera phylogeny available (Song et al., 2020). All of the following processes were performed in the R programming environment. First, an evaluation of taxon overlap between the Song et al., 2020 phylogeny and the used taxon sampling was conducted and a tip substitution based on the most-restrictive taxonomic rank was implemented. Species that had no match in the taxon sampling were pruned from the tree using drop.tip in ape (Paradis and Schliep, 2019). 12 species that had an unequivocal sister group (the same genus or family/subfamily branch only had one tip) were manually added with addTip in TreeTools (Smith, 2019). This function had the advantage that not only the new edge length could be defined but also the edge length of the already present sister group as well. To keep the tree ultrametric the edge length of the new tip was randomized between 0 and the edge length of the sister group. These procedures lead to a subset of 153 taxa (out of 343 taxa) which were used for phylogenetic comparative statistics.

To check if the phylogenetic signal was influenced by adding taxa manually to the tree, 1000 trees with random new edge lengths for the added taxa were calculated. Phylogenetic signal, in the form of Kmult (the multivariate version of Blomberg’s K (Blomberg et al., 2003) for the polynomial coefficients of all 1000 randomized trees was calculated with physignal() in geomorph with 999 iterations (Adams and Otárola-Castillo, 2013). The tree with a phylogenetic signal closest to the mean K of all randomized trees was used in all further statistical analyses.

### Allometric and phylogenetic signal correction

Potential correlation between the bite efficiency profile in form of polynomial coefficients and the log mandible length was done jointly while estimating phylogenetic effects using a phylogenetically informed (phylogenetic generalized least squares [PGLS]) regression. The function procD.lm.pgls (Geomorph: (Adams and Otárola-Castillo, 2013), was performed with 10.000 permutations. The PGLS regression implementation used assumes a Brownian Motion evolutionary model. For further analysis of the phylogenetic and allometric corrected data, the residuals of the PGLS function were used.

To ascertain which evolutionary model fits the data best the package mvMorph (Clavel et al., 2015, v. 1.1.6) was used. The function mvgls() uses a maximum likelihood approach (method = “LL” in function) to fit multivariate linear models to multivariate data. The data was tested against the evolutionary models of Brownian Motion, Ornstein-Uhlenbeck and Early Burst for the predictors of diet and family group. The best fit was determined using the AICc criterion.

### Statistics

The mandible length for all statistical tests was measured as the distance between the in-lever line and the mandible tip point. All further tests were done in the R programming environment. To explore the patterns of variation within the bite efficiency on the gnathal edge a Principal Component Analysis (PCA) on the polynomial coefficients was performed in R using the function gm.prcomp in geomorph.

One-Way-ANOVAs were done to test for potential correlations between biological factors and the proximal as well as a distal mechanical advantage using the function aov() in stats (R Core Team, 2022). Afterwards, a Tukey’s range was performed using TukeyHSD() to determine which diet groups were significantly different to each other. Correlation between the bite efficiency profile (polynomial coefficients) and the factors diet, mandible length and family grouping was tested with a factorial ANOVA using procD.lm() in geomorph.

## Results

All statistical analyses were conducted with the mechanical advantage of the adductor and abductor mandible muscle. All following results, if not explicitly stated are reported for the adductor muscle. Results concerning the abductor muscle are reported in the supplements.

### Bite efficiency profile patterns

A polynomial curve of the 9th order had the best fit based on the AIC scores. The biting efficiency profile (BEP) of Orthoptera had the highest MA at the proximal end, which varied between 0.94 and 0.63 at the closer muscle and 0.35 to 0.09 at the opener muscle. It then decreased by ∼70% along the biting area until it reached its lowest point at the most distal end. The Outgroups in contrast showed a lower proximal MA which ranged from 0.65 to 0.48 at the closer and 0.4 to 0.16 at the opener muscle. The progression of the MA was visually similar between most orthopteran superfamilies (Fig. 2) with some showing areas of decreased negative slope. The Gryllotalpoidea (mole and ant crickets), Grylloidea (crickets), Hagloidea (grigs) and Tanaoceroidea (desert long-horned grasshoppers) had this slope at the 25% position and the Tetrigoidea (pygmy grasshoppers) at the 50% position. Superfamilies with a more uniform BEP were the Schizodactyloidea (dune crickets), Stenopelmatoidea, Rhaphidophoroidea (cave crickets), and Tettigonioidea.

**Fig. 2:**
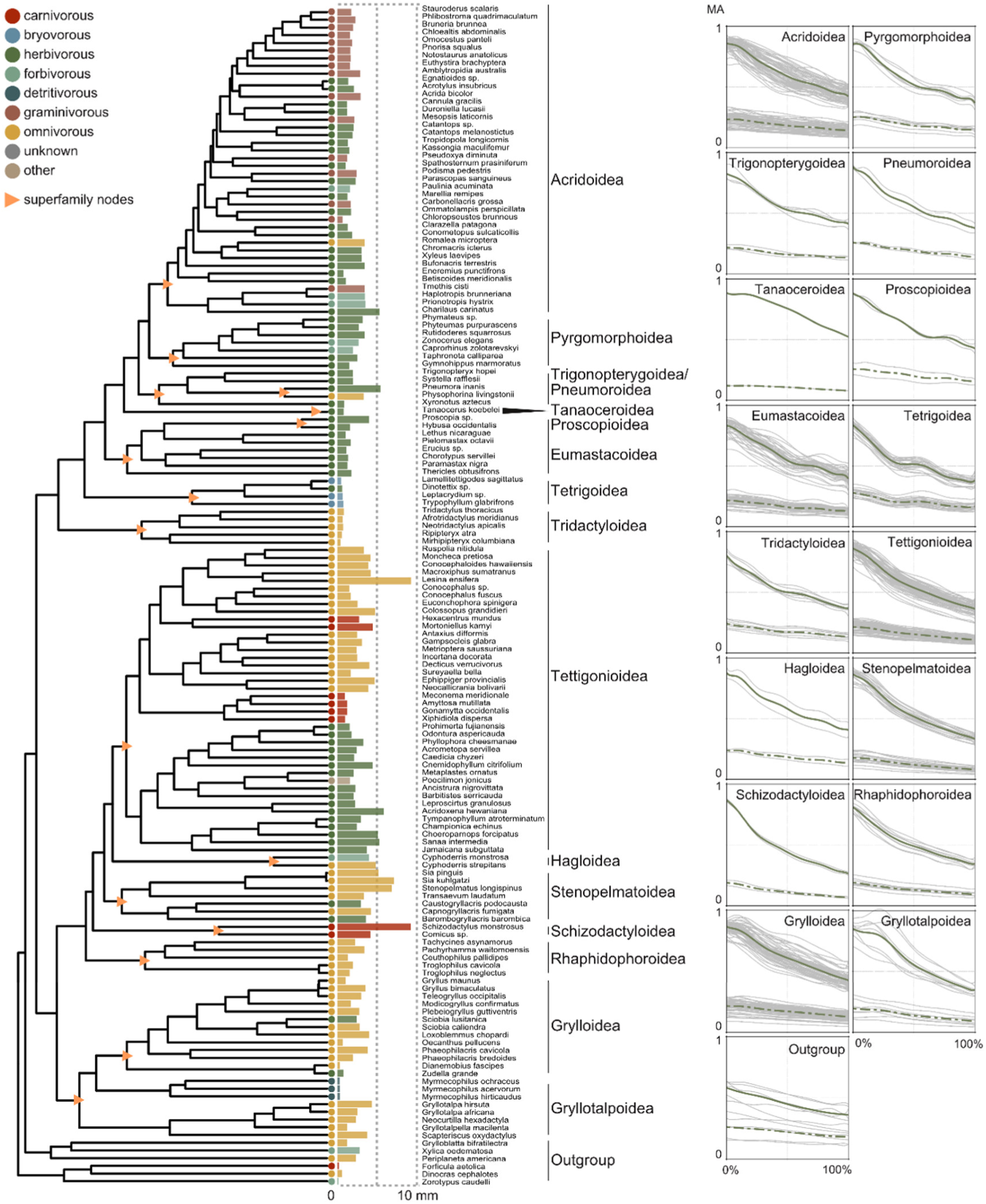
The evolution of dietary categories, mandible size and mechanical advantage in Orthoptera. Left: Phylogenetic relationships (adapted from Song et al. (2020)) with coloured bars indicating dietary category and absolute mandible length in mm. Right: Mean mechanical advantage (green lines) and its variation among species of each superfamily (grey lines; whole dataset; N=345) for the closer muscle (solid line) and the opener muscle (dotted line). X-axis = Percentage of gnathal edge with 0% as the most proximal and 100% as the most distal point of the gnathal edge.

### Allometric and phylogenetic signal

There was a significant (P<0.05) phylogenetic signal in the BEP as well as the distal MA, proximal MA and the MA difference between proximal and distal MA (from now on only referred to as MA difference) (Tab. 2).Kmult had values below 1 for the BEP as well as the distal MA and MA difference and hence a lower resemblance than expected under Brownian Motion (BM). The proximal MA showed a Kmult over 1 which means a higher resemblance of closely related taxa than expected under BM. The comparison of evolutionary models for the polynomial coefficients with family and diet showed that the Brownian motion evolutionary model was the preferred model for both factors (S 7). Because mandible length in the dataset had a 62-fold size variation from 0.18 to 9.29 mm (mean 2.18 mm) correlation between BEPs and size was tested. An ANOVA comparing the log-transformed mandible length of the whole data set to the BEPs showed significant (P<0.05) but very weak influences of mandible length on the BEP variation (R2 = 0.11) (S 6). PGLS regression of BEPs on log mandible length again had significant but very low explanatory values (R2 = 0.086, P<0.05).

**Tab. 2:** Results of phylogenetic signal testing. Numbers show Kmult with all results being significant.

| data | muscle | $K_{mult}$ | Z | P |
| --- | --- | --- | --- | --- |
| polynomial coefficients | closer | 0.97 | 15.20 | 0.001 |
| polynomial coefficients | opener | 0.35 | 3.91 | 0.001 |
| MA distal | closer | 0.61 | 5.61 | 0.001 |
| MA proximal |  | 2.50 | 17.31 | 0.001 |
| MA difference |  | 1.73 | 15.78 | 0.001 |
| MA distal | opener | 0.51 | 4.60 | 0.001 |
| MA proximal |  | 0.34 | 2.32 | 0.01 |
| MA difference |  | 0.38 | 2.87 | 0.002 |

### Exploring the morphospace occupation of the biting efficiency profile

A principal component analysis of the polynomial coefficients showed that PCs 1 and 2 (Fig. 3, S 1, S 2) described 81% of the variation in the data set which increased after allometric/phylogenetic correction by 4% (S 4). No distinct functional, ecological or taxonomical groups, except the outgroups, were determined based on visual inspection. The morphospace occupation of each species was mostly explained by distal MA and MA difference. Both factors increased from the lower left to the upper right quadrant. Gryllotalpoidea showed a higher disparity within the PCA than other Orthoptera with a recognizable distinction between the two families of Gryllotalpidae (mole crickets) and the Myrmecophilidae (ant-crickets). The ant crickets clustered around the central point of the dataset whereas the Gryllotalpidae separated into the lower right quadrant. Gryllotalpidae also showed the highest disparity among the superfamilies (S 10) and they were, besides the outgroups, the only superfamily with a significant difference in disparity compared to all other superfamilies. Disparity analysis of the diet groups gave a high disparity in the omnivore guild with significant differences between graminivorous and mixed-herbivorous (S 9). They also had a higher distribution of species in the PCA analysis after phylogenetic and allometric correction (S 1).

**Fig. 3:**
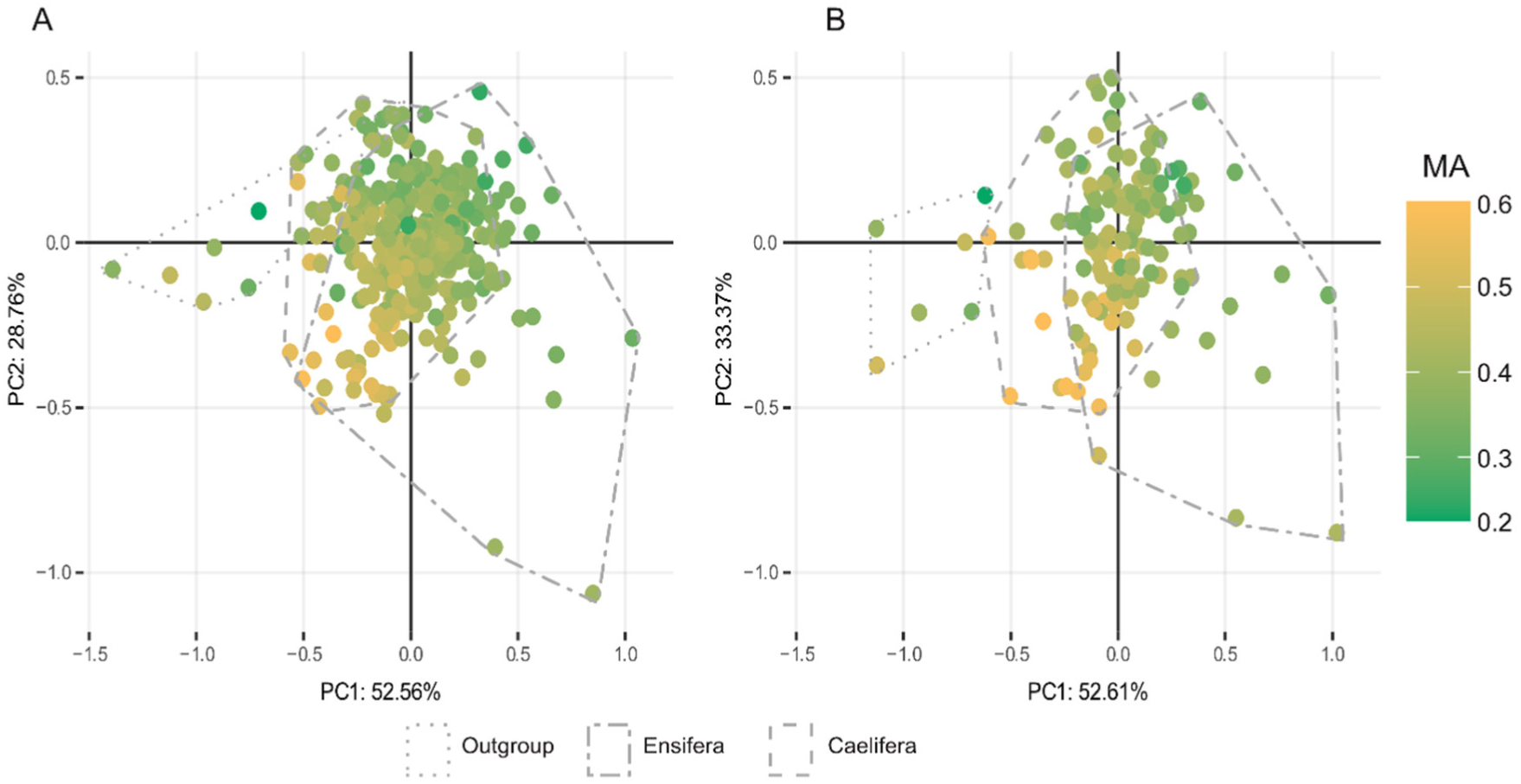
The mechanical advantage space of Orthoptera. Shown here are principal components 1 and 2 with (A) the complete taxon set (N = 345) and (B) the subset with phylogenetic and allometric correction (N = 153). Distal MAs are shown with a colour coding from yellow to green.

### Interaction of BEP, MA, diet and taxonomic grouping

All statistical tests were run with the whole taxon set (N = 343) as well as the subset (N = 153) represented in the phylogenetic tree. Results from all tests will be reported in brackets with the first number representing the whole taxon set and the second representing the subset. The phylogenetic and allometric corrected results were restricted to the subset. To make values easier to compare increases and decreases will be reported compared to the subset.

Multifactorial ANOVAs with the coefficients of the BEP as the dependent variables and the point MA measurements (distal MA, proximal MA and MA difference), family groups and diet groups as the independent variable, displayed a significant correlation before and after allometric/phylogenetic correction in all tests (Tab. 4).

**Tab. 3:**
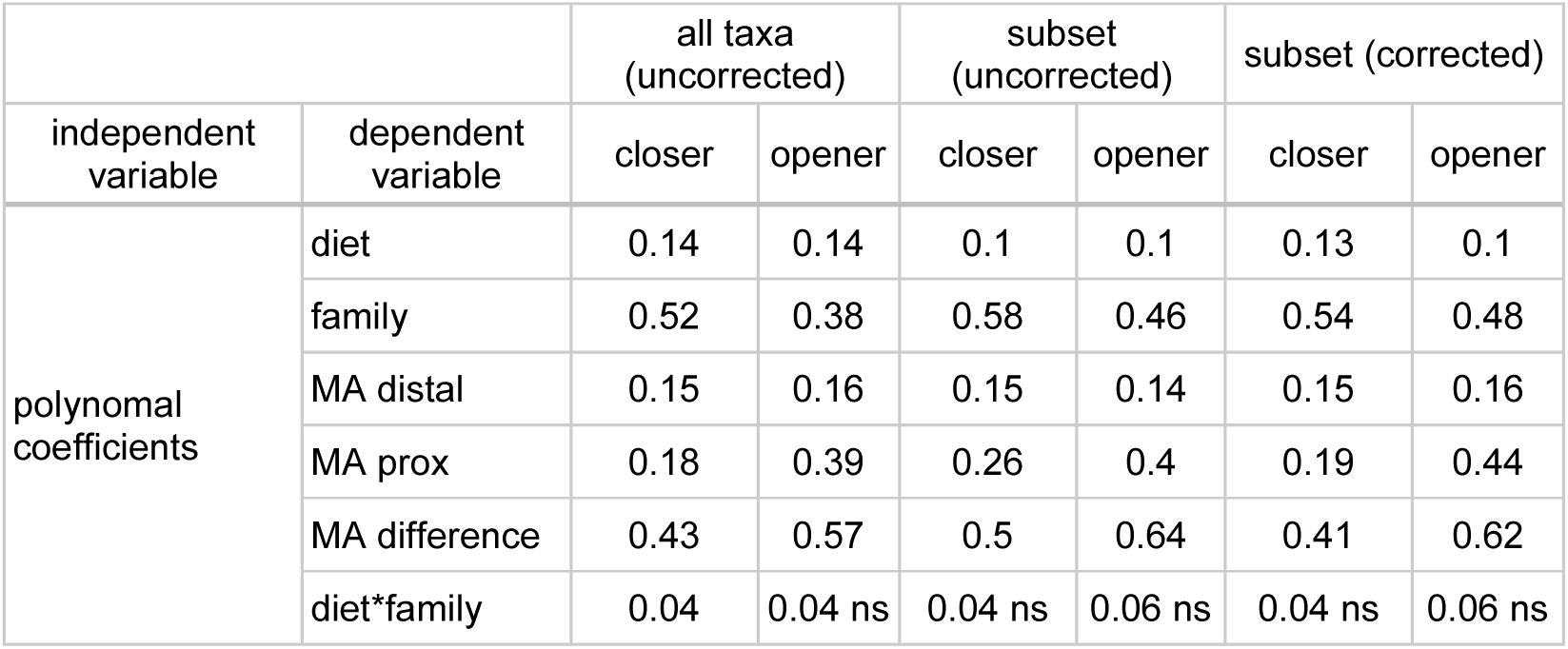
R2 values of One-Way ANOVA and MANOVA. Correlation tests were done between distal and proximal MA to diet and family groups as well as interaction terms between diet and family. Subset means the phylogenetic subset present in the phylogenetic tree (N = 153). For the allometric and phylogenetic corrected test, PGLS-corrected residuals were used in the ANOVA. ns = non-significant. * marks tested interaction in MANOVA

**Tab. 4:**
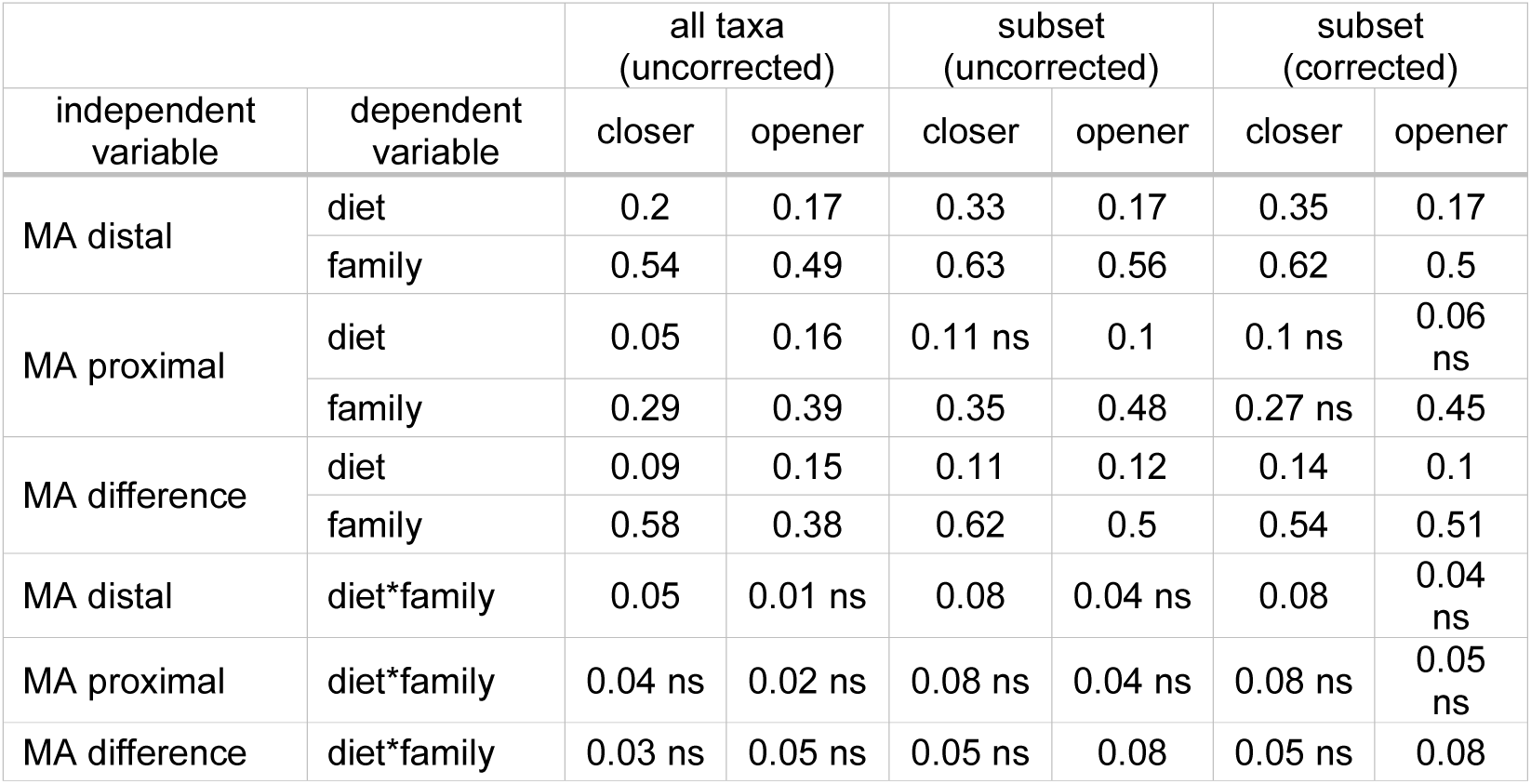
R2 values of multifactorial ANOVA and MANOVA between the polynomial coefficients and diet as well as family groupings. Subset means the phylogenetic subset present in the phylogenetic tree (N = 153). For the allometric and phylogenetic corrected test, PGLS-corrected residuals were used in the ANOVA. ns = non-significant. * marks tested interaction in MANOVA

Diet had low explanatory power for BEP variance (R2 = 0.14; R2 = 0.1) compared to taxonomic family grouping (R2 = 0.54; R2 = 0.58). After taking phylogenetic and allometric non-independence under consideration an increase in correlation to diet (R2 = 0.13) and a decrease in correlation to family grouping was found. Distal MA (R2 = 0.15; R2 = 0.15) and proximal MA (R2 = 0.18; R2 = 0.26) both had low correlation to the BEP and both values stayed the same after phylogenetic and allometric correction. The MA difference had a higher explanatory power of the variance in the BEP (R2 = 0.43; R2 = 0.5) which decreased with the removal of the allometric and phylogenetic signal (R2 = 0.47).

Because the single point values of proximal MA and distal MA showed a significant correlation to the BEP, potential correlations to diet and family were also tested (Tab. 3). A significant correlation between diet and the distal MA was found (R2 = 0.20; R2 = 0.33) which increased with correction for size and phylogenetic relationship (R2 = 0.35). The family grouping had a much higher explanatory power for distal MA (R2 = 0.54; R2 = 0.63) which only slightly decreased after accounting for phylogeny (R2 = 0.62). The proximal MA showed a significant correlation to diet in the whole taxon set (R2 = 0.05) which was lost when in the subset even before correction. Correlation between family grouping and proximal MA was significant in both taxon sets (R2 = 0.29; R2 = 0.35) but not after allometric/phylogenetic correction. MA difference again showed a significant but low correlation to diet (R2 = 0.09; R2 = 0.011) which increased after correction (R2 = 0.14) and a higher correlation to family grouping (R2 = 0.58; R2 = 0.62) which slightly decreased after correction (R2 = 0.54). Interaction between diet and family resulted in no significant correlation to either distal MA, proximal MA or MA difference.

### Exploration of the MA variation in diet guilds

The explanatory power of diet for the BEP and the point MA measurements was in all statistical tests low but significant in all tests. To explore the variation between the different diet guilds a post hoc Tukey’s test of the One-Ways-ANOVAs was performed (Fig. 4, S 8). It showed that the mean of the proximal MA was not significantly different between any of the dietary guilds. The mean of the distal MA and the MA difference only had a significant difference between some diet guilds. Detritivorous and carnivorous Orthoptera showed the lowest distal MA and graminivorous the highest. Significant differences were detected between the carnivorous guild and all other guilds except the detritivorous one, the “others” and the bryovorous one. In addition, a significant difference was detected between the bryovorous guild and the mixed-herbivorous as well as the graminivorous guild. The MA difference only showed statistical differences between carnivorous and the sole phytophagous groups of forbivorous, graminivorous and mixed-herbivorous). Omnivorous and graminivorous diet guilds also were significantly different from each other.

**Fig. 4:**
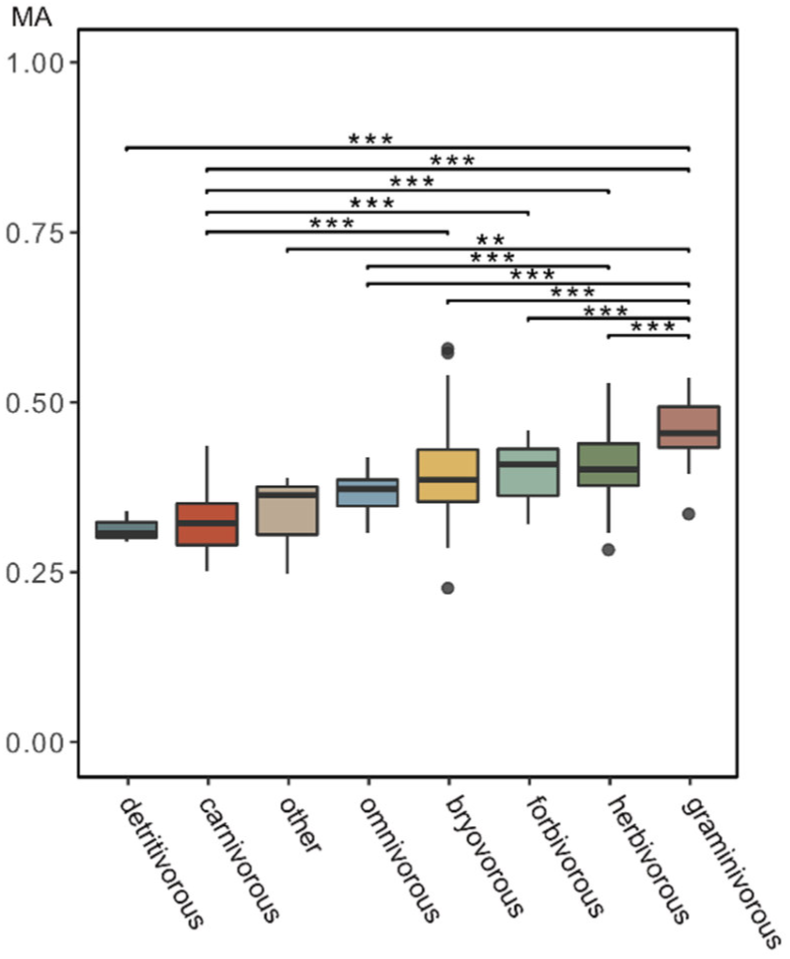
Boxplots of the distal mechanical advantage of each dietary category with significant differences (after Tukey’s Range Test) shown above the plots. *** = P < 0.001; ** = P < 0.01.

## Discussion

For many taxa, it was shown that mouthpart phenotypes correlate with aspects of the feeding ecology (Blomberg et al., 2003; Evans et al., 2007; Kienle and Berta, 2016; Meyers et al., 2006; Montaña and Winemiller, 2013; Nogueira et al., 2009; Prevosti et al., 2012; Santana et al., 2022) which even lead to predictions of feeding preferences (Firmat et al., 2010). Due to the involvement of mouthparts in the cutting and mastication of food, an adaptation to increase food consumption efficiency generally is expected (Stephens and Krebs, 2019). In insects, an increased intake rate determines survival not only by the acquisition of nutrients. Feeding is also dangerous because it can increase the risk of predation (Bernays, 1997). Contrary to the established notion of a tight correlation of mandible shape with feeding ecology, our results show that diet is not the main determinant behind the diversification of bite force transmission efficiency and, consequently, mandible shape in Orthoptera. Instead, phylogeny drives and/or constrains this diversification. Bite efficiency progression varies more with family grouping than with dietary grouping and together with a strong phylogenetic signal indicates that, overall, closely related species are less similar than expected under a Brownian motion model of evolution. This is in contrast to qualitative studies based on the visual sorting of mandible types into different dietary categories (Gangwere, 1966, 1965; Isely, 1944; Patterson, 1984; Wheater and Evans, 1989) and studies for other animal groups. In Mesozoic mammals shorter jaws and increased MAs are associated with herbivory whereas intermediate and longer jaws and lower MAs are associated with insectivory and carnivory(Morales-García et al., 2021). Similar results were found for rodents (Anderson et al., 2014; Missagia et al., 2021), birds (Olsen, 2017) and reptiles (Bestwick et al., 2021).

In Orthoptera, despite an enormous mandible length disparity of a factor of 100, correlation to MA variation along the gnathal edge held very low explanatory power. As long as the ratio between the in-lever and out-lever keeps within a steady range, no shape adaptation to diet or size is necessary. A similar effect can be observed in the suction feeding in teleost fishes where size differences of multiple traits (gape width, buccal length, cross-sectional area of the epaxial muscle, lengths of the epaxialis, and buccal cavity moment arms) can result in a similar performance (Wainwright et al., 2007).

Another possibility is that muscle volume and angle as well as muscle physiology compensate for a less-than-optimal mandible shape which might render shape adaptations redundant. Bernays (Bernays, 1986) showed that grass-feeding grasshoppers have an increased head mass due to an increase in mandible musculature which might be enough to increase food throughput to meet nutritional needs. Muscle physiology was linked to foraging habits in ants (Gronenberg et al., 1997) where the composition of muscle fibre types determined mandible speed. However, such changes in muscle mass were small given the different dietary categories across Orthoptera analysed here. In evolutionary terms, a change in mandible shape, e.g., along the molar region appears to be a simpler adaptation than larger changes in muscle angles, masses and, consequently, associated changes of other head parts to accommodate such different muscle configurations.

### A closer look at gnathal edge disparity in Orthoptera

Despite the overall low explanatory power of diet for mandible shape, we also found that the closer muscle had the highest distal MA within grass feeders and the lowest in detritivores and carnivorous Orthoptera. This followed the expectation that tougher plant matter like grasses necessitates a higher force transmission and a lower MA is more advantageous in carnivory, due to an increase in biting speed. Despite this, the explanatory power of diet for the MA was low and no distinct dietary functional groups for the mechanical advantage profile along the gnathal edge could be determined. What little correlation could be found was mostly defined by differences between graminivorous and carnivorous Orthoptera to the other dietary guilds. Grass-feeding is a mechanically demanding process with a higher chance of material wear due to silica deposits and tougher venation in grasses (Raupp, 1985; Wright and Vincent, 1996). This suggests that adaptation pressure to the diet increases with mechanical challenges but otherwise a multi-tool approach is sufficient.

Despite the generally low explanatory power of feeding categories for mandible shape and efficiency, there are some notable examples of specialization in our dataset. The superfamily group with the highest disparity are the Gryllotalpoidae which includes the Gryllotalpidae (mole-crickets) and Myrmecophilidae (ant-crickets). They are two families with very specialized and different living environments. The ant crickets live inquiline with ants and obtain food by either grooming or trophallaxis for which they have modified mouthparts (Parmentier, 2020; Komatsu and Maruyama, 2016; Wetterer and Hugel, 2008). Mole crickets are adapted for digging and live mostly underground. They have a nearly prognathic head orientation with a shortened-out lever distance and an increase in the MA in the proximal part of the mandible. The genus Oecanthus similarly shows a more prognathic head orientation (Hudson, 1945) and shows similar angling of the mandible and a slight increase in mechanical advantage. A case of dietary conservatism on a family level can be found in the Tetrigidae (Kuřavová et al., 2017) which prefer algae, mosses and lychees. Their mandible shows a specialized scraping ridge at their molar area (Kuřavová et al., 2014; Kuravova and Kocarek, 2016), which was theorized to follow convergent adaptation in algae grazers (Arens, 1994; Kuřavová et al., 2017).

In summary, although there are many shape differences observable for orthopteran mandibles, our results show that there are also many intermediate mandible forms which likely lead to the observed very low explanatory values. On the other hand, mandibles shapes can largely differ although the dietary category is the same. For example, the mandibles of the forb-feeding Acrididae Paulinia acuminata look different from the forb-feeding Prophalangopsidae Cyphoderris monstrosa.

### Omnivory: A comfortable spot

While Orthoptera are generally perceived as phytophagous animals, that mostly feed on plant matters most Orthoptera cannot be categorized into only one dietary group. Many species feed indiscriminately on plant matter but do not refuse protein sources like dead arthropods when provided (Bernays and Chapman, 1970; Ingrisch and Köhler, 1998). Even strictly defined grass-feeders like the Gomphocerinae or Romalea microptera can be observed to feed on other arthropods (Clark, 1948; Richardson et al., 2012). The advantage of an omnivorous approach is that animals always have access to sustenance in form of abundant plant matter as well as highly nutritious protein-rich food sources like animal matter.

Our results are in line with this facultative omnivory of many species. Orthopteran mandibles operate in a performative state with multiple peaks but rather a plateau in which morphological disparity can move around without changing the performance optimum. This is a characteristic of morphological stasis in which environments may change dramatically but morphology does not follow (Wake et al., 1983). Similarly, Zelditch (Zelditch et al., 2020) found that while tree squirrels are morphologically specialized they are ecological generalists which leads to a persistent optimum. The beaks of Darwin finches were equally found to lack diet specialisation in preference for versatility when defined on a macroevolutionary scale (Navalón et al., 2019).

### End statement

Orthoptera originated ∼355 Million years ago (Song et al., 2020) and persisted despite massive environmental and climatic changes. However, one major factor of their microhabitat, plants, remained relatively stable in the sense that food was always highly abundant. Radiations took place mostly in the Mesozoic within Ensifera in correlation with the angiosperm radiation and for the Caelifera during the Cenozoic coinciding with the spreading of grasslands (Song et al., 2020) but the principle food source always remained in high abundance. Such stable microhabitat environments with abundant nutritional access possibly have led to a lack of specification and diversification pressure in mouthpart configuration.

## Author contributions

CE: conceptualization, data curation, formal analysis, investigation, methodology, project administration, visualization, writing-original draft, writing-review and editing

TvdK: investigation, methodology, resources, software, writing-review and editing

TF: investigation, methodology, resources, software, writing-review and editing

JUH: investigation, methodology, resources, software, writing-review and editing

FW: investigation, methodology, resources, software, writing-review and editing

PTR: investigation, writing-review and editing

MF: investigation, writing-review and editing

AB: conceptualization, funding acquisition, investigation, project administration, resources, supervision, writing-review and editing

## Acknowledgements

Loaning of museum specimens where kindly organized by B. Price (NHMUK), H. Zettel (NHMV), D. Zimmermann (NHMV), J. Deckert (MfN), R. Peters (ZFMK) and L. Hendrich (ZSM). We also thank the member of the KIT, DESY for access to their facilities and their rendered assistance.

## Supplementary material

*S 1: PC1 and 2 of the polynomial coefficients. Diet groups used in colour coding. A/B: Complete taxon set at the closer (A) and opener (B) muscle without corrections (N = 345). C/D: Phylogenetic subset at the closer (C) and opener (D) muscle after phylogenetic and allometric correction (N = 153)*.

*S 2: PC1 and 2 of the polynomial coefficients. Superfamily groups are used in colour coding. A/B: Complete taxon set at the closer (A) and opener (B) muscle without corrections (N = 345). C/D: Phylogenetic subset at the closer (C) and opener (D) muscle after phylogenetic and allometric correction (N = 153)*.

*S 3: Render of mandible 3D Models done in Blender. All Mandibles are scaled to have the same Volume. Colours represent diet preference according to the literature*.

*S 4: Explanation of variance for each PCA axis (in %) of the whole uncorrected taxon set and the phylogenetic and allometric corrected subset. Included are the results for the opener and closer muscle*.

*S 5: R 2 - Results of the PGLS regression between subsets polynomial coefficients, MA distal, MA proximal and MA difference as the independent variable and the log of mandible length as dependent variable. ns = not significant*

*S 6: R 2 - R 2 - Results of the testing for allometric signal on the whole taxon set. the polynomial coefficients, MA distal, MA proximal and MA difference as the independent variable and the log of mandible length as depenent variable. ns = not significant*

*S 7: AIC and log likelihood scores (LL) from evolutionary model fit test for the polynomial coefficients as a function of diet and family grouping. Tested evolutionary models: BM = Brownian Motion; OU = Ornstein-Uhlenbeck; EB = Early burst*.

*S 8: Posthoc Tukey Test Results for distal MA, proximal MA and MA difference*.

*S 9: Results of testing for disparity within diet groups for polynomial coefficients, distal MA, proximal, MA and MA difference on the closer and opener muscle Groups with significant difference of mean markes with *** = P<0.001; ** = P <0.01; P < 0.005*

*S 10: Results of testing for disparity within diet superfamily groups for polynomial coefficients, distal MA, proximal, MA and MA difference on the closer and opener muscle Groups with significant difference of mean markes with *** = P<0.001; ** = P <0.01; P < 0.005*

*S 11: List of Species used in Analysis and acquired data*

*S 12: List with Literature Results for diet preference*

## References

Adams, D.C., Otárola-Castillo, E., 2013. geomorph: an r package for the collection and analysis of geometric morphometric shape data. Methods Ecol. Evol. 4, 393–399. https://doi.org/10.1111/2041-210X.12035

Aguirre-Segura, A., Arcos, M., Moyano, F., Pascual, F., 1987. Tipos adaptivos de morfologia mandibular en algunas espesies de ortópteros ibéricos. Graellsia XLIII, 225–235.

Anderson, P.S., Renaud, S., Rayfield, E.J., 2014. Adaptive plasticity in the mouse mandible. BMC Evol. Biol. 14, 85. https://doi.org/10.1186/1471-2148-14-85

Arens, W., 1994. Striking convergence in the mouthpart evolution of stream-living algae grazers. J. Zool. Syst. Evol. Res. 32, 319–343. https://doi.org/10.1111/j.1439-0469.1994.tb00490.x

Behmer, S.T., 2009. Insect Herbivore Nutrient Regulation. Annu. Rev. Entomol. 54, 165–187. https://doi.org/10.1146/annurev.ento.54.110807.090537

Bennack, D.E., 1981. The effects of mandible morphology and photosynthetic pathway on selective herbivory in grasshoppers. Oecologia 51, 281–283. https://doi.org/10.1007/BF00540615

Bernays, E.A., 1997. Feeding by lepidopteran larvae is dangerous. Ecol. Entomol. 22, 121–123. https://doi.org/10.1046/j.1365-2311.1997.00042.x

Bernays, E.A., 1986. Diet-Induced Head Allometry among Foliage-Chewing Insects and its Importance for Graminivores. Sci. New Ser. 231, 495–497.

Bernays, E.A., Chapman, R.F., 1970. Food Selection by Chorthippus parallelus (Zetterstedt) (Orthoptera: Acrididae) in the Field. J. Anim. Ecol. 39, 383–394. https://doi.org/10.2307/2977

Bernays, E.A., Jarzembowski, E.A., Malcolm, S.B., 1991. Evolution of Insect Morphology in Relation to Plants [and Discussion]. Philos. Trans. Biol. Sci. 333, 257–264.

Berthaume, M.A., Winchester, J., Kupczik, K., 2019. Ambient occlusion and PCV (portion de ciel visible): A new dental topographic metric and proxy of morphological wear resistance. PLOS ONE 14, e0215436. https://doi.org/10.1371/journal.pone.0215436

Bestwick, J., Unwin, D.M., Henderson, D.M., Purnell, M.A., 2021. Dental microwear texture analysis along reptile tooth rows: complex variation with non-dietary variables. R. Soc. Open Sci. 8, 201754. https://doi.org/10.1098/rsos.201754

Blomberg, S.P., Garland JR. T., Ives, A.R., 2003. Testing for Phylogenetic Signal in Comparative Data: Behavioral Traits Are More Labile. Evolution 57, 717–745. https://doi.org/10.1111/j.0014-3820.2003.tb00285.x

Bozdogan, H., 1987. Model selection and Akaike’s Information Criterion (AIC): The general theory and its analytical extensions. Psychometrika 52, 345–370. https://doi.org/10.1007/BF02294361

Burkhardt, F., Smith, S., Kohn, D., Montgomery, W., 1985. A calendar of the correspondence of Charles Darwin, 1821-1882. J. Hist. Biol. 18.

Carroll, A.M., Wainwright, P.C., Huskey, S.H., Collar, D.C., Turingan, R.G., 2004. Morphology predicts suction feeding performance in centrarchid fishes. J. Exp. Biol. 207, 3873–3881. https://doi.org/10.1242/jeb.01227

Chapman, R.F., 1995. Mechanics of Food Handling by Chewing Insects, in: Chapman, R.F., de Boer, G. (Eds.), Regulatory Mechanisms in Insect Feeding. Springer US, Boston, MA, pp. 3–31. https://doi.org/10.1007/978-1-4615-1775-7_1

Clark, E.J., 1948. Studies in the Ecology of British Grasshoppers. Trans. R. Entomol. Soc. Lond. 99, 173–222. https://doi.org/10.1111/j.1365-2311.1948.tb01235.x

Clavel, J., Escarguel, G., Merceron, G., 2015. mvmorph: an r package for fitting multivariate evolutionary models to morphometric data. Methods Ecol. Evol. 6, 1311–1319. https://doi.org/10.1111/2041-210X.12420

Clifton, K.B., Motta, P.J., 1998. Feeding Morphology, Diet, and Ecomorphological Relationships among Five Caribbean Labrids (Teleostei, Labridae). Copeia 1998, 953–966. https://doi.org/10.2307/1447342

Clissold, F., 2007. The Biomechanics of Chewing and Plant Fracture: Mechanisms and Implications. Adv. Insect Physiol. 34, 317–372. https://doi.org/10.1016/S0065-2806(07)34006-X

Clissold, F.J., Sanson, G.D., Read, J., Simpson, S.J., 2009. Gross vs. net income: How plant toughness affects performance of an insect herbivore. Ecology 90, 3393–3405. https://doi.org/10.1890/09-0130.1

Corbin, C.E., Lowenberger, L.K., Gray, B.L., 2015. Linkage and trade-off in trophic morphology and behavioural performance of birds. Funct. Ecol. 29, 808–815. https://doi.org/10.1111/1365-2435.12385

Edgecombe, G., Richter, S., Wilson, G., 2003. The mandibular gnathal edges: homologous structures across Mandibulata? Afr. Invertebr. 44, 115–135.

ElEla, S.A., ElSayed, W., Nakamura, K., 2010. Mandibular structure, gut contents analysis and feeding group of orthopteran species collected from different habitats of Satoyama area within Kanazawa City, Japan. J. Threat. Taxa 849–857. https://doi.org/10.11609/JoTT.o2346.849-57

Evans, A.R., Wilson, G.P., Fortelius, M., Jernvall, J., 2007. High-level similarity of dentitions in carnivorans and rodents. Nature 445, 78–81. https://doi.org/10.1038/nature05433

Feilich, K.L., López-Fernández, H., 2019. When Does Form Reflect Function? Acknowledging and Supporting Ecomorphological Assumptions. Integr. Comp. Biol. 59, 358–370. https://doi.org/10.1093/icb/icz070

Firmat, C., Rodrigues, H.G., Renaud, S., Hutterer, R., Garcia-Talavera, F., Michaux, J., 2010. Mandible morphology, dental microwear, and diet of the extinct giant rats Canariomys (Rodentia: Murinae) of the Canary Islands (Spain). Biol. J. Linn. Soc. 101, 28–40. https://doi.org/10.1111/j.1095-8312.2010.01488.x

Gangwere, S.K., 1966. Relationships between the Mandibles, Feeding Behavior, and Damage Inflicted on Plants by the Feeding of Certain Acridids (Orthoptera) 1, 5.

Gangwere, S.K., 1965. The structural adaptations of mouthparts in Orthoptera and Allis.

Gangwere, S.K., 1961. A monograph on food selection in Orthoptera. Trans. Am. Entomol. Soc. 1890–87, 67–230.

Gangwere, S.K., McKinney, J.C., Ernemann, M.A., Bland, R.G., 1998. Food Selection and Feeding Behavior in Selected Acridoidea (Insecta: Orthoptera) of the Canary Islands, Spain. J. Orthoptera Res. 1–21. https://doi.org/10.2307/3503485

Gangwere, S.K., Spiller, D.O., 1995. Food Selection and Feeding Behavior in Selected Orthoptera sen. lat. of the Balearic Islands, Spain. J. Orthoptera Res. 147–160. https://doi.org/10.2307/3503470

Gronenberg, W., Paul, J., Just, S., Hölldobler, B., 1997. Mandible muscle fibers in ants: fast or powerful? Cell Tissue Res. 289, 347–361. https://doi.org/10.1007/s004410050882

Grossnickle, D.M., 2020. Feeding ecology has a stronger evolutionary influence on functional morphology than on body mass in mammals. Evolution 74, 610–628. https://doi.org/10.1111/evo.13929

Harzing, A.W., 2007. Publish or Perish.

Hess, R., 2010. Blender Foundations: The Essential Guide to Learning Blender 2.6. Focal Press.

Hillerton, J.E., Reynolds, S.E., Vincent, J.F.V., 1982. On the Indentation Hardness of Insect Cuticle. J. Exp. Biol. 96, 45–52. https://doi.org/10.1242/jeb.96.1.45

Hochuli, D.F., 2001. Insect herbivory and ontogeny: How do growth and development influence feeding behaviour, morphology and host use?: ONTOGENETIC CONSTRAINTS ON INSECT HERBIVORY. Austral Ecol. 26, 563–570. https://doi.org/10.1046/j.1442-9993.2001.01135.x

Hochuli, D.F., 1996. The Ecology of Plant/insect Interactions: Implications of Digestive Strategy for Feeding by Phytophagous Insects. Oikos 75, 133. https://doi.org/10.2307/3546331

Hudson, G.B., 1945. A study of the tentorium in some orthopteroid Hexapoda. J. Entomol. Soc. South. Afr. 8, 71–90. https://doi.org/10.10520/AJA00128789_3869

Ingrisch, S., Köhler, G., 1998. Die Heuschrecken Mitteleuropas. Westarp Wissenschaften.

Isely, F.B., 1944. Correlation Between Mandibular Morphology And Food Specificity In Grasshoppers1. Ann. Entomol. Soc. Am. 37, 47–67. https://doi.org/10.1093/aesa/37.1.47

Kaltenbach, A., 1990. The predatory Saginae. Tettigoniidae Biol. Syst. Evol. 1, 280–320.

Kang, L., Gan, Y., Li, S., 1999. The Structural Adaptation of Mandibles and Food Specificity in Grasshoppers on Inner Mongolian Grasslands. J. Orthoptera Res. 257. https://doi.org/10.2307/3503442

Kaufmann, T., 1965. Biological Studies on Some Bavarian Acridoidea (Orthoptera), with Special Reference to Their Feeding Habits. Ann. Entomol. Soc. Am. 58, 791–801. https://doi.org/10.1093/aesa/58.6.791

Khokhriakov, I., Beckmann, F., Lottermoser, L., 2017. Integrated control system environment for high-throughput tomography (No. PUBDB-2018-01600), Proceedings of SPIE. Presented at the Developments in X-Ray Tomography XI, SPIE. https://doi.org/10.1117/12.2275863

Kienle, S.S., Berta, A., 2016. The better to eat you with: the comparative feeding morphology of phocid seals (Pinnipedia, Phocidae). J. Anat. 228, 396–413. https://doi.org/10.1111/joa.12410

Komatsu, T., Maruyama, M., 2016. Taxonomic recovery of the ant cricket Myrmecophilus albicinctus from M. americanus (Orthoptera, Myrmecophilidae). ZooKeys 97–106. https://doi.org/10.3897/zookeys.589.7739

Kuřavová, K., Hajduková, L., Kočárek, P., 2014. Age-related mandible abrasion in the groundhopper Tetrix tenuicornis (Tetrigidae, Orthoptera). Arthropod Struct. Dev. 43, 187–192. https://doi.org/10.1016/j.asd.2014.02.002

Kuravova, K., Kocarek, P., 2016. Mandibular morphology and dietary preferences in two pygmy molecrickets of the genus Xya (Orthoptera: Tridactylidae). Turk. J. Zool. 40, 720–728. https://doi.org/10.3906/zoo-1510-19

Kuřavová, K., Šipoš, J., Wahab, Rodzaj.A., Kahar, R.S., Kočárek, P., 2017. Feeding patterns in tropical groundhoppers (Tetrigidae): a case of phylogenetic dietary conservatism in a basal group of Caelifera. Zool. J. Linn. Soc. 179, 291–302. https://doi.org/10.1111/zoj.12474

Le Gall, M., Behmer, S.T., 2014. Effects of Protein and Carbohydrate on an Insect Herbivore: The Vista from a Fitness Landscape. Integr. Comp. Biol. 54, 942–954. https://doi.org/10.1093/icb/icu102

Lupu, G., 2007. Carnivorous and omnivorous species of Orthoptera order recorded in the Danube Delta Biosphere Reserve. Sci. Ann. DDI.

Metzger, K.A., Herrel, A., 2005. Correlations between lizard cranial shape and diet: a quantitative, phylogenetically informed analysis. Biol. J. Linn. Soc. 86, 433–466.

Meyers, J.J., Herrel, A., Nishikawa, K.C., 2006. Morphological correlates of ant eating in horned lizards (Phrynosoma). Biol. J. Linn. Soc. 89, 13–24. https://doi.org/10.1111/j.1095-8312.2006.00654.x

Missagia, R.V., Patterson, B.D., Krentzel, D., Perini, F.A., 2021. Insectivory leads to functional convergence in a group of Neotropical rodents. J. Evol. Biol. 34, 391–402. https://doi.org/10.1111/jeb.13748

Montaña, C.G., Winemiller, K.O., 2013. Evolutionary convergence in Neotropical cichlids and Nearctic centrarchids: evidence from morphology, diet, and stable isotope analysis. Biol. J. Linn. Soc. 109, 146–164. https://doi.org/10.1111/bij.12021

Moosmann, J., Ershov, A., Weinhardt, V., Baumbach, T., Prasad, M.S., LaBonne, C., Xiao, X., Kashef, J., Hofmann, R., 2014. Time-lapse X-ray phase-contrast microtomography for in vivo imaging and analysis of morphogenesis. Nat. Protoc. 9, 294–304. https://doi.org/10.1038/nprot.2014.033

Morales-García, N.M., Gill, P.G., Janis, C.M., Rayfield, E.J., 2021. Jaw shape and mechanical advantage are indicative of diet in Mesozoic mammals. Commun. Biol. 4, 1–14. https://doi.org/10.1038/s42003-021-01757-3

Morris, P.J.R., Cobb, S.N.F., Cox, P.G., 2018. Convergent evolution in the Euarchontoglires. Biol. Lett. 14, 20180366. https://doi.org/10.1098/rsbl.2018.0366

Navalón, G., Bright, J.A., Marugán-Lobón, J., Rayfield, E.J., 2019. The evolutionary relationship among beak shape, mechanical advantage, and feeding ecology in modern birds*. Evolution 73, 422–435. https://doi.org/10.1111/evo.13655

Nogueira, M.R., Peracchi, A.L., Monteiro, L.R., 2009. Morphological correlates of bite force and diet in the skull and mandible of phyllostomid bats. Funct. Ecol. 23, 715–723. https://doi.org/10.1111/j.1365-2435.2009.01549.x

Olsen, A.M., 2017. Feeding ecology is the primary driver of beak shape diversification in waterfowl. Funct. Ecol. 31, 1985–1995. https://doi.org/10.1111/1365-2435.12890

Orthoptera Species File. Version 5.0/5.0. [WWW Document], 2021. URL http://Orthoptera.SpeciesFile.org (accessed 4.13.21).

Otte, D., Joern, A., 1976. On Feeding Patterns in Desert Grasshoppers and the Evolution of Specialized Diets. Proc. Acad. Nat. Sci. Phila. 128, 89–126.

Paradis, E., Schliep, K., 2019. ape 5.0: an environment for modern phylogenetics and evolutionary analyses in R. Bioinformatics 35, 526–528.

Patterson, B.D., 1984. Correlation Between Mandibular Morphology and Specific Diet of Some Desert Grassland Acrididae (Orthoptera). Am. Midl. Nat. 111, 296–303. https://doi.org/10.2307/2425324

Patterson, B.D., 1983. Grasshopper Mandibles and the Niche Variation Hypothesis. Evolution 37, 375–388. https://doi.org/10.2307/2408345

Powell, P.L., Roy, R.R., Kanim, P., Bello, M.A., Edgerton, V.R., 1984. Predictability of skeletal muscle tension from architectural determinations in guinea pig hindlimbs. J. Appl. Physiol. 57, 1715– 1721. https://doi.org/10.1152/jappl.1984.57.6.1715

Prevosti, F.J., Turazzini, G.F., Ercoli, M.D., Hingst-Zaher, E., 2012. Mandible shape in marsupial and placental carnivorous mammals: a morphological comparative study using geometric morphometrics. Zool. J. Linn. Soc. 164, 836–855.

Püffel, F., Pouget, A., Liu, X., Zuber, M., van de Kamp, T., Roces, F., Labonte, D., 2021. Morphological determinants of bite force capacity in insects: a biomechanical analysis of polymorphic leaf-cutter ants. J. R. Soc. Interface 18, 20210424. https://doi.org/10.1098/rsif.2021.0424

R Core Team, 2022. R: A Language and Environment for Statistical Computing. R Foundation for Statistical Computing, Vienna, Austria.

Raupp, M.J., 1985. Effects of leaf toughness on mandibular wear of the leaf beetle, Plagiodera versicolora. Ecol. Entomol. 10, 73–79. https://doi.org/10.1111/j.1365-2311.1985.tb00536.x

Richardson, M.L., Reagel, P.F., Mitchell, R.F., Whitman, D.W., 2012. Opportunistic Carnivory by Romalea microptera (Orthoptera: Acrididae). Ann. Entomol. Soc. Am. 105, 28–35. https://doi.org/10.1603/AN11057

Richter, S., Edgecombe, G.D., Wilson, G.D.F., 2002. The lacinia mobilis and Similar Structures – a Valuable Character in Arthropod Phylogenetics? Zool. Anz. - J. Comp. Zool. 241, 339–361. https://doi.org/10.1078/0044-5231-00083

Rühr, P.T., Blanke, A., 2022. forceX and forceR: A mobile setup and r package to measure and analyse a wide range of animal closing forces. Methods Ecol. Evol. 13, 1938–1948. https://doi.org/10.1111/2041-210X.13909

Rühr, P.T., van de Kamp, T., Faragó, T., Hammel, J.U., Wilde, F., Borisova, E., Edel, C., Frenzel, M., Baumbach, T., Blanke, A., 2021. Juvenile ecology drives adult morphology in two insect orders. Proc. R. Soc. B Biol. Sci. 288, 20210616. https://doi.org/10.1098/rspb.2021.0616

Samways, M.J., Osborn, R., Saunders, T.L., 1997. Mandible Form Relative to the Main Food Type in Ladybirds (Coleoptera: Coccinellidae). Biocontrol Sci. Technol. 7, 275–286. https://doi.org/10.1080/09583159730974

Sanson, G., 2006. The biomechanics of browsing and grazing. Am. J. Bot. 93, 1531–1545. https://doi.org/10.3732/ajb.93.10.1531

Santana, S.E., Grossnickle, D.M., Sadier, A., Patterson, E., Sears, K.E., 2022. Bat Dentitions: A Model System for Studies at the Interface of Development, Biomechanics, and Evolution. Integr. Comp. Biol. 62, 762–773. https://doi.org/10.1093/icb/icac042

Schindelin, J., Arganda-Carreras, I., Frise, E., Kaynig, V., Longair, M., Pietzsch, T., Preibisch, S., Rueden, C., Saalfeld, S., Schmid, B., Tinevez, J.-Y., White, D.J., Hartenstein, V., Eliceiri, K., Tomancak, P., Cardona, A., 2012. Fiji: an open-source platform for biological-image analysis. Nat. Methods 9, 676–682. https://doi.org/10.1038/nmeth.2019

Schofield, R.M.S., Nesson, M.H., Richardson, K.A., 2002. Tooth hardness increases with zinc-content in mandibles of young adult leaf-cutter ants. Naturwissenschaften 89, 579–583. https://doi.org/10.1007/s00114-002-0381-4

Schoonhoven, L.M., Van Loon, J.J., Dicke, M., 2005. Insect-plant biology. Oxford University Press on Demand.

Smith, M.R., 2019. TreeTools: create, modify and analyse phylogenetic trees. Comprehensive R Archive Network. https://doi.org/10.5281/zenodo.3522725

Smith, T.R., Capinera, J.L., 2005. Mandibular Morphology of Some Floridian Grasshoppers (Orthoptera: Acrididae). Fla. Entomol. 88, 204–207.

Snodgrass, R., 1935. Principles of Insect Morphology. McGraw-Hill Publishing Co., New York.

Song, H., Béthoux, O., Shin, S., Donath, A., Letsch, H., Liu, S., McKenna, D.D., Meng, G., Misof, B., Podsiadlowski, L., Zhou, X., Wipfler, B., Simon, S., 2020. Phylogenomic analysis sheds light on the evolutionary pathways towards acoustic communication in Orthoptera. Nat. Commun. 11, 4939. https://doi.org/10.1038/s41467-020-18739-4

Stampanoni, M., Groso, A., Isenegger, A., Mikuljan, G., Chen, Q., Bertrand, A., Henein, S., Betemps, R., Frommherz, U., Böhler, P., Meister, D., Lange, M., Abela, R., 2006. Trends in synchrotron-based tomographic imaging: the SLS experience, in: Developments in X-Ray Tomography V. Presented at the Developments in X-Ray Tomography V, SPIE, pp. 193–206. https://doi.org/10.1117/12.679497

Stayton, C.T., 2006. Testing Hypotheses of Convergence with Multivariate Data: Morphological and Functional Convergence Among Herbivorous Lizards. Evolution 60, 824–841. https://doi.org/10.1111/j.0014-3820.2006.tb01160.x

Stephens, D.W., Krebs, J.R., 2019. Foraging Theory, Foraging Theory. Princeton University Press. https://doi.org/10.1515/9780691206790

Tan, M.K., Artchwakom, T., Wahab, R.A., Lee, C.-Y., Belabut, D.M., Tan, H.T.W., 2017. Overlooked flower-visiting Orthoptera in Southeast Asia. J. Orthoptera Res. 26, 143–153. https://doi.org/10.3897/jor.26.15021

Uvarov, S.B., 1966. Grasshoppers and locusts. A handbook of general acridology. Volume I. Anatomy, physiology, development, phase polymorphism, introduction to taxonomy. Anti-Locust Res. Cent. Camb. Univ. Press 1.

Vincent, J.F.V., Wegst, U.G.K., 2004. Design and mechanical properties of insect cuticle. Arthropod Struct. Dev. 33, 187–199. https://doi.org/10.1016/j.asd.2004.05.006

Wainwright, P., Carroll, A.M., Collar, D.C., Day, S.W., Higham, T.E., Holzman, R.A., 2007. Suction feeding mechanics, performance, and diversity in fishes. Integr. Comp. Biol. 47, 96–106. https://doi.org/10.1093/icb/icm032

Wainwright, P.C., Richard, B.A., 1995. Predicting patterns of prey use from morphology of fishes. Environ. Biol. Fishes 44, 97–113. https://doi.org/10.1007/BF00005909

Wake, D.B., Roth, G., Wake, M.H., 1983. On the problem of stasis in organismal evolution. J. Theor. Biol. 101, 211–224. https://doi.org/10.1016/0022-5193(83)90335-1

Weihmann, T., Reinhardt, L., Weißing, K., Siebert, T., Wipfler, B., 2015. Fast and Powerful: Biomechanics and Bite Forces of the Mandibles in the American Cockroach Periplaneta americana. PLoS ONE 10, e0141226. https://doi.org/10.1371/journal.pone.0141226

Westneat, M.W., 2004. Evolution of Levers and Linkages in the Feeding Mechanisms of Fishes1. Integr. Comp. Biol. 44, 378–389. https://doi.org/10.1093/icb/44.5.378

Wetterer, J., Hugel, S., 2008. Worldwide Spread of the Ant Cricket Myrmecophilus americanus, a Symbiont of the Longhorn Crazy Ant, Paratrechina longicornis. Sociobiology 52, 157–165.

Wheater, C.P., Evans, M.E.G., 1989. The mandibular forces and pressures of some predacious Coleoptera. J. Insect Physiol. 35, 815–820. https://doi.org/10.1016/0022-1910(89)90096-6

Wilde, F., Ogurreck, M., Greving, I., Hammel, J.U., Beckmann, F., Hipp, A., Lottermoser, L., Khokhriakov, I., Lytaev, P., Dose, T., Burmester, H., Müller, M., Schreyer, A., 2016. Micro-CT at the imaging beamline P05 at PETRA III. AIP Conf. Proc. 1741, 030035. https://doi.org/10.1063/1.4952858

Wright, W., Vincent, J.F.V., 1996. Herbivory and the mechanics of fracture in plants. Biol. Rev. Camb. Philos. Soc. U. K.

Yushkevich, P.A., Piven, J., Hazlett, H.C., Smith, R.G., Ho, S., Gee, J.C., Gerig, G., 2006. User-guided 3D active contour segmentation of anatomical structures: Significantly improved efficiency and reliability. NeuroImage 31, 1116–1128. https://doi.org/10.1016/j.neuroimage.2006.01.015

Zelditch, M.L., Li, J., Swiderski, D.L., 2020. Stasis of functionally versatile specialists. Evolution 74, 1356–1377. https://doi.org/10.1111/evo.13956

