## Supplement S1 to S9 for "Bite force transmission and mandible shape in grasshoppers, crickets, and allies is largely dependent on phylogeny, not diet"

*S 1: PC1 and 2 of the polynomial coefficients. Diet groups used in colour coding. A/B: Complete taxon set at the closer (A) and opener (B) muscle without corrections (N = 345). C/D: Phylogenetic subset at the closer (C) and opener (D) muscle after phylogenetic and allometric correction (N = 153).*

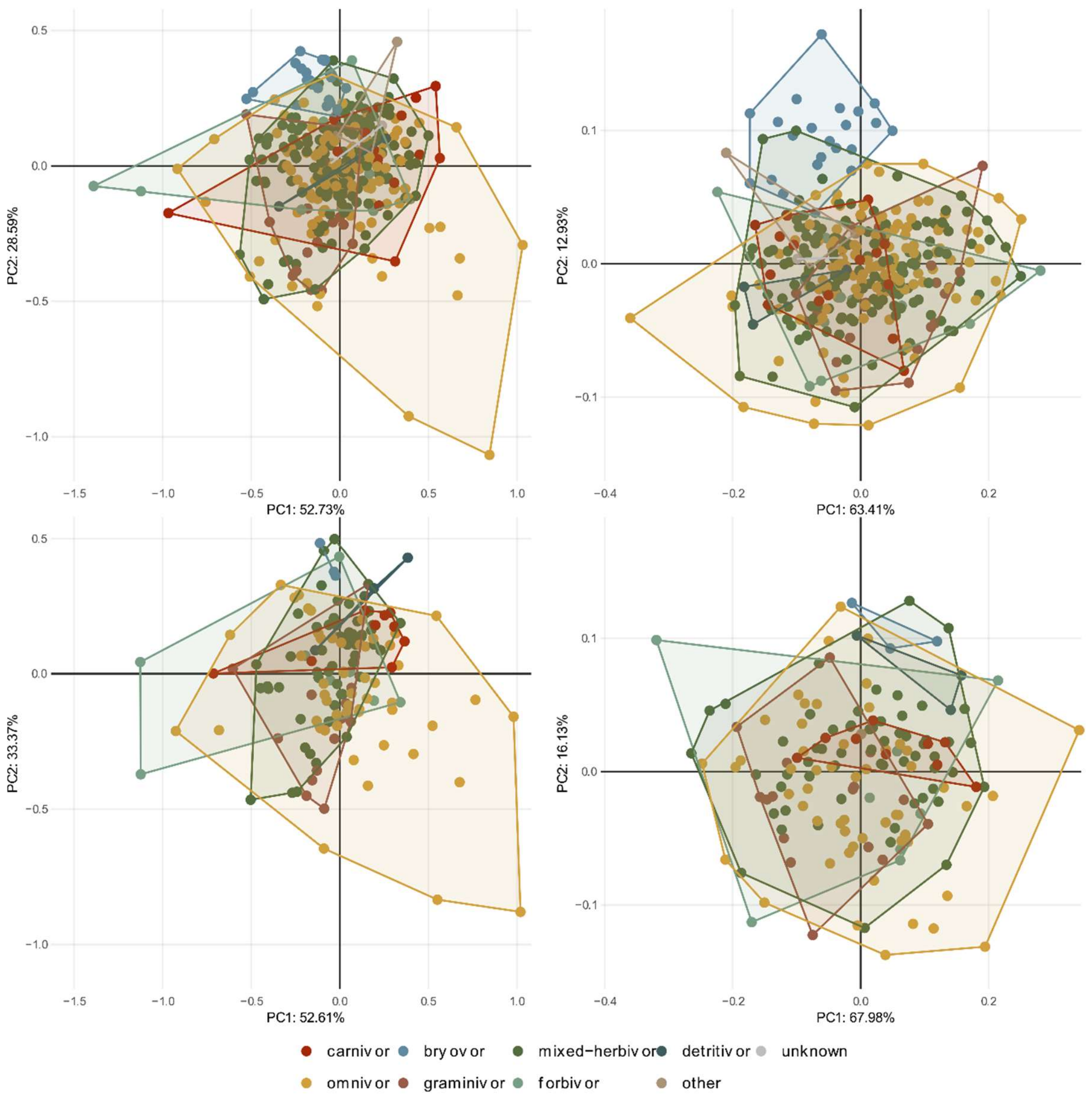

*S 2: PC1 and 2 of the polynomial coefficients. Superfamily groups are used in colour coding. A/B: Complete taxon set at the closer (A) and opener (B) muscle without corrections (N = 345). C/D: Phylogenetic subset at the closer (C) and opener (D) muscle after phylogenetic and allometric correction (N = 153).*

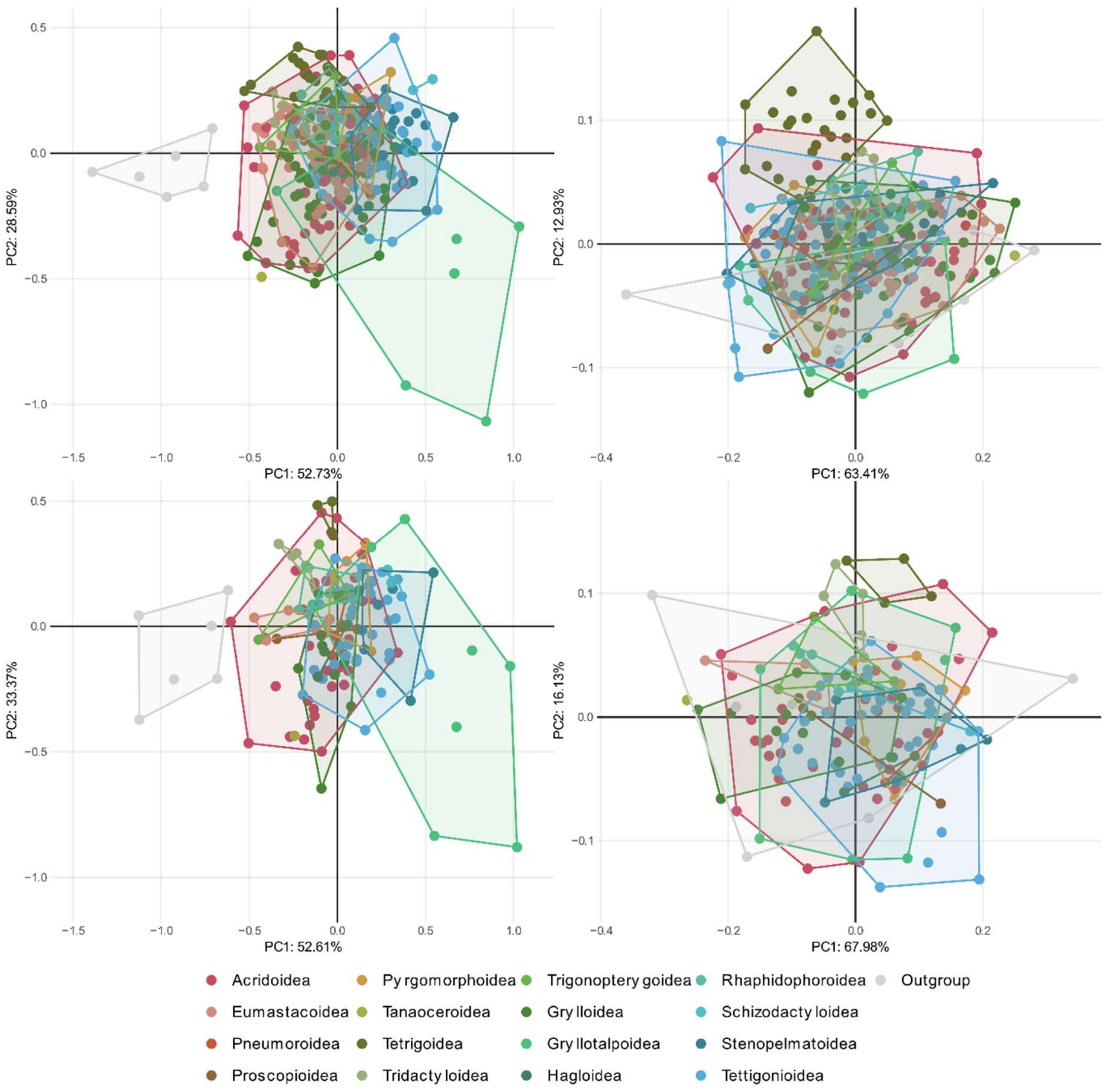

*S 3: Render of mandible 3D Models done in Blender. All Mandibles are scaled to have the same Volume.*

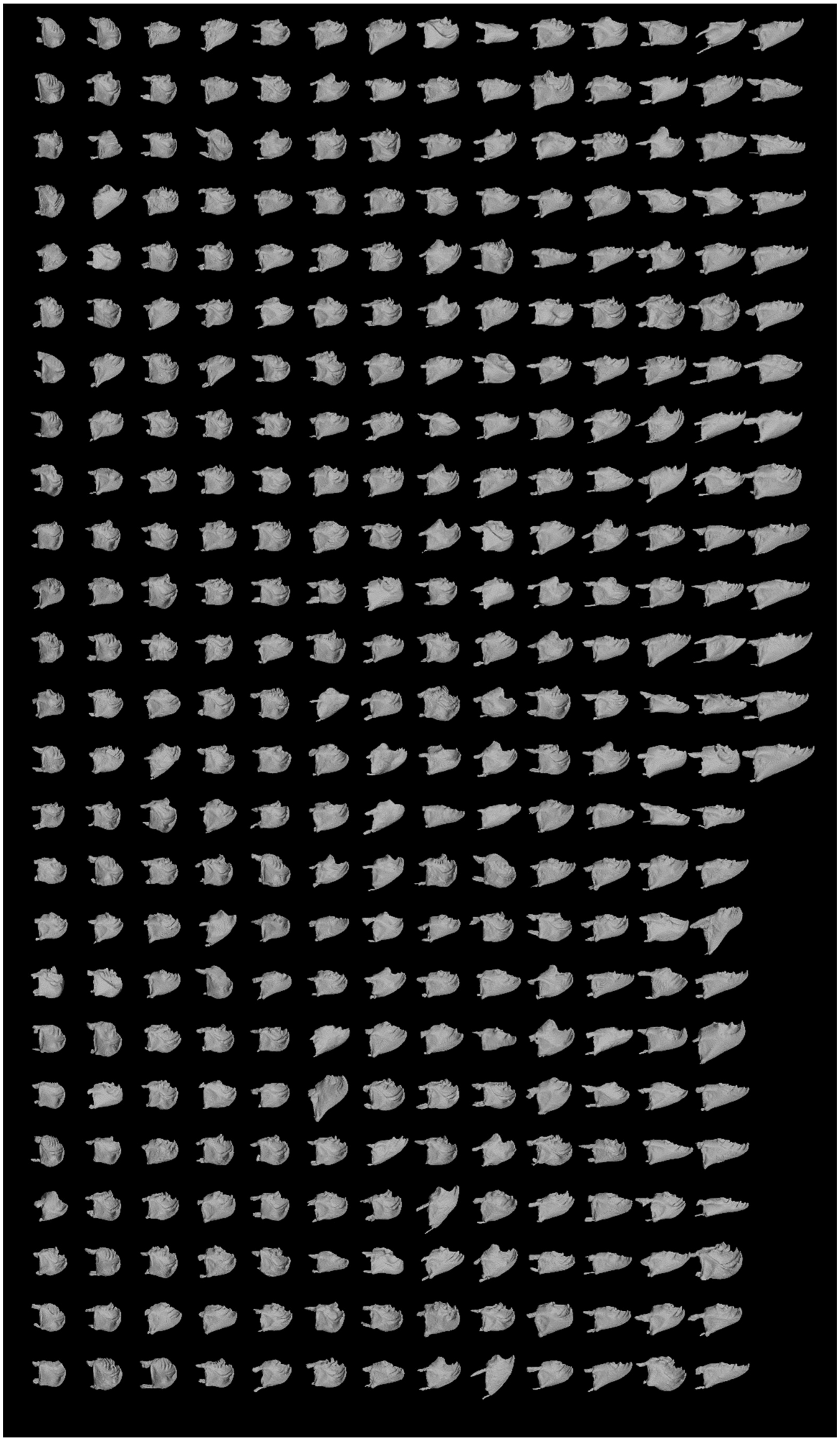

*S 4: Explanation of variance for each PCA axis (in %) of the whole uncorrected taxon set and the phylogenetic and allometric corrected subset. Included are the results for the opener and closer muscle.*

| | all taxa (uncorrected) | | subset (uncorrected) | | subset (corrected) | |
|---|---|---|---|---|---|---|
| axis | closer | opener | closer | opener | closer | opener |
| PC1 | 52.73 | 63.41 | 59.13 | 70.42 | 52.61 | 67.98 |
| PC2 | 28.59 | 12.93 | 28.11 | 11.87 | 33.37 | 16.13 |
| PC3 | 7.69 | 9.98 | 4.42 | 7.21 | 4.99 | 5.95 |
| PC4 | 3.79 | 6.05 | 2.68 | 4.19 | 2.94 | 4.09 |
| PC5 | 2.67 | 2.74 | 2.23 | 2.13 | 2.25 | 2.02 |
| PC6 | 2.03 | 1.92 | 1.54 | 1.63 | 1.66 | 1.41 |
| PC7 | 1.04 | 1.58 | 0.75 | 1.35 | 0.82 | 1.27 |
| PC8 | 0.9 | 0.92 | 0.72 | 0.79 | 0.79 | 0.73 |
| PC9 | 0.46 | 0.36 | 0.36 | 0.33 | 0.52 | 0.33 |
| PC10 | 0.09 | 0.11 | 0.06 | 0.09 | 0.07 | 0.09 |

*S 5: $R^2$ - Results of the PGLS regression between subsets polynomial coefficients, MA distal, MA proximal and MA difference as the independent variable and the log of mandible length as dependent variable. ns = not significant*

| | closer | opener |
|---|---|---|
| polynomial coefficient | 0.09 | 0.05 |
| MA distal | 0.01 ns | 0.07 |
| MA proximal | 0.07 | 0.11 |
| MA difference | 0.04 | 0.01 ns |

*S 6: $R^2$ - Results of the testing for allometric signal on the whole taxon set.  the polynomial coefficients, MA distal, MA proximal and MA difference as the independent variable and the log of mandible length as depenent variable. ns = not significant*

| | closer | opener |
|---|---|---|
| polynomial coefficients | 0.11 | 0.02 |
| MA distal | 0.03 | 0.07 |
| MA proximal | 0.18 | 0.13 |
| MA difference | 0.08 | 0 ns |

*S 7: AIC and log likelihood scores (LL) from evolutionary model fit test for the polynomial coefficients as a function of diet and family grouping. Tested evolutionary models: BM = Brownian Motion; OU = Ornstein-Uhlenbeck; EB = Early burst.*

| | muscle | AIC_BM | AIC_OU | AIC_EB | AIC_λ | LL_BM | LL_OU | LL_EB | LL_lambda |
|---|---|---|---|---|---|---|---|---|---|
| diet | closer | -1798.33 | -1992.16 | -1798.33 | -2071.05 | 1798.33 | 1992.16 | 1798.33 | 2071.05 |
| diet | opener | -3399.55 | -3647.51 | -3399.55 | -3703.85 | 3399.55 | 3647.51 | 3399.55 | 3703.85 |
| family | closer | -1487.15 | -1753.24 | -1487.15 | -1763.12 | 1487.15 | 1753.24 | 1487.15 | 1763.12 |
| family | opener | -2789.31 | -3045.16 | -2789.31 | -3060.05 | 2789.31 | 3045.16 | 2789.31 | 3060.05 |

*S 8: Posthoc Tukey Test Results for distal MA, proximal MA and MA difference. Groups with significant difference of mean markes with *** = P<0.001; ** = P <0.01; P < 0.005*

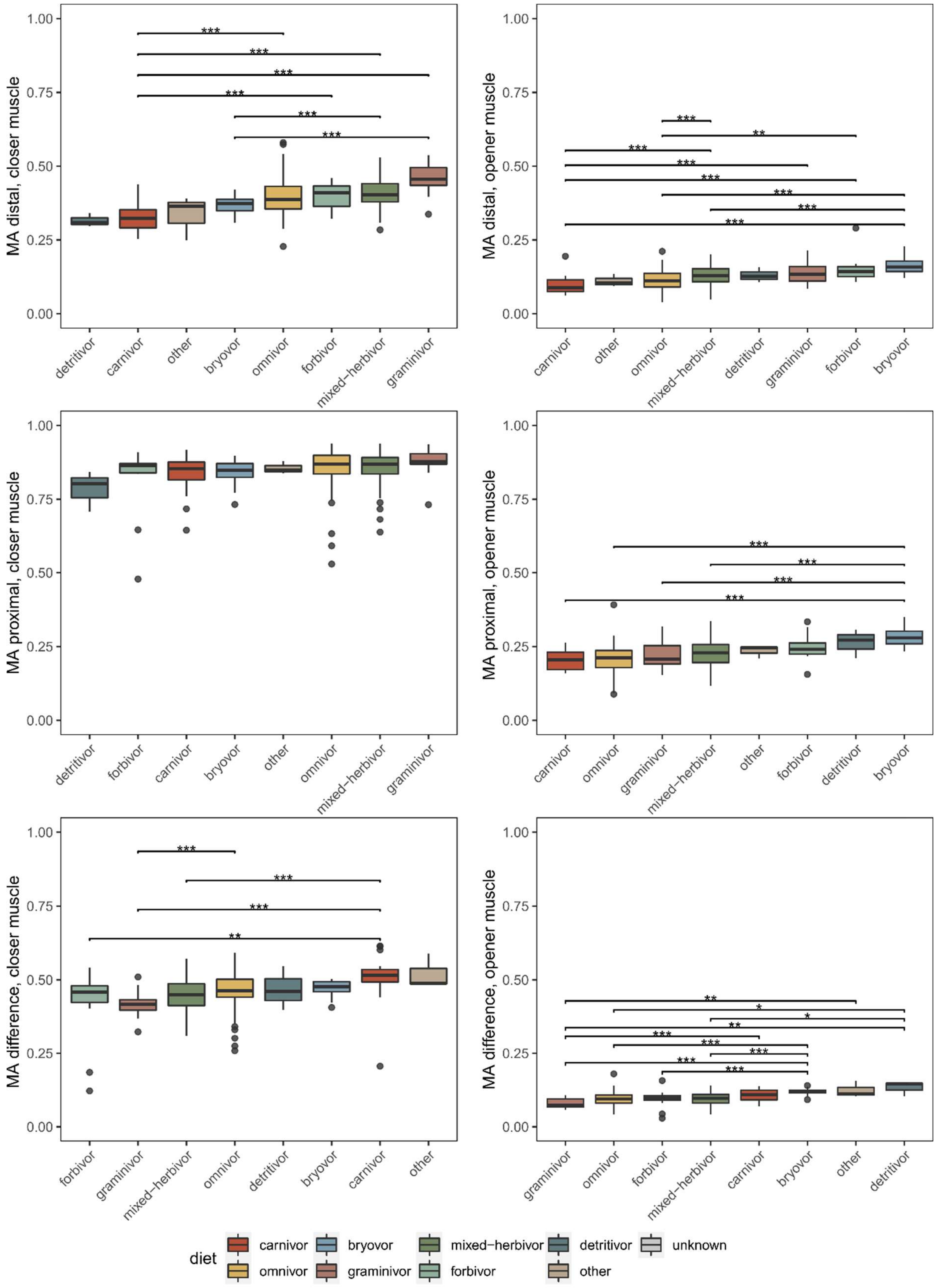

*S 8: Results of testing for disparity within diet groups for polynomial coefficients, distal MA, proximal, MA and MA difference on the closer and opener muscle Groups with significant difference of mean markes with *** = P<0.001; ** = P <0.01; P < 0.005*

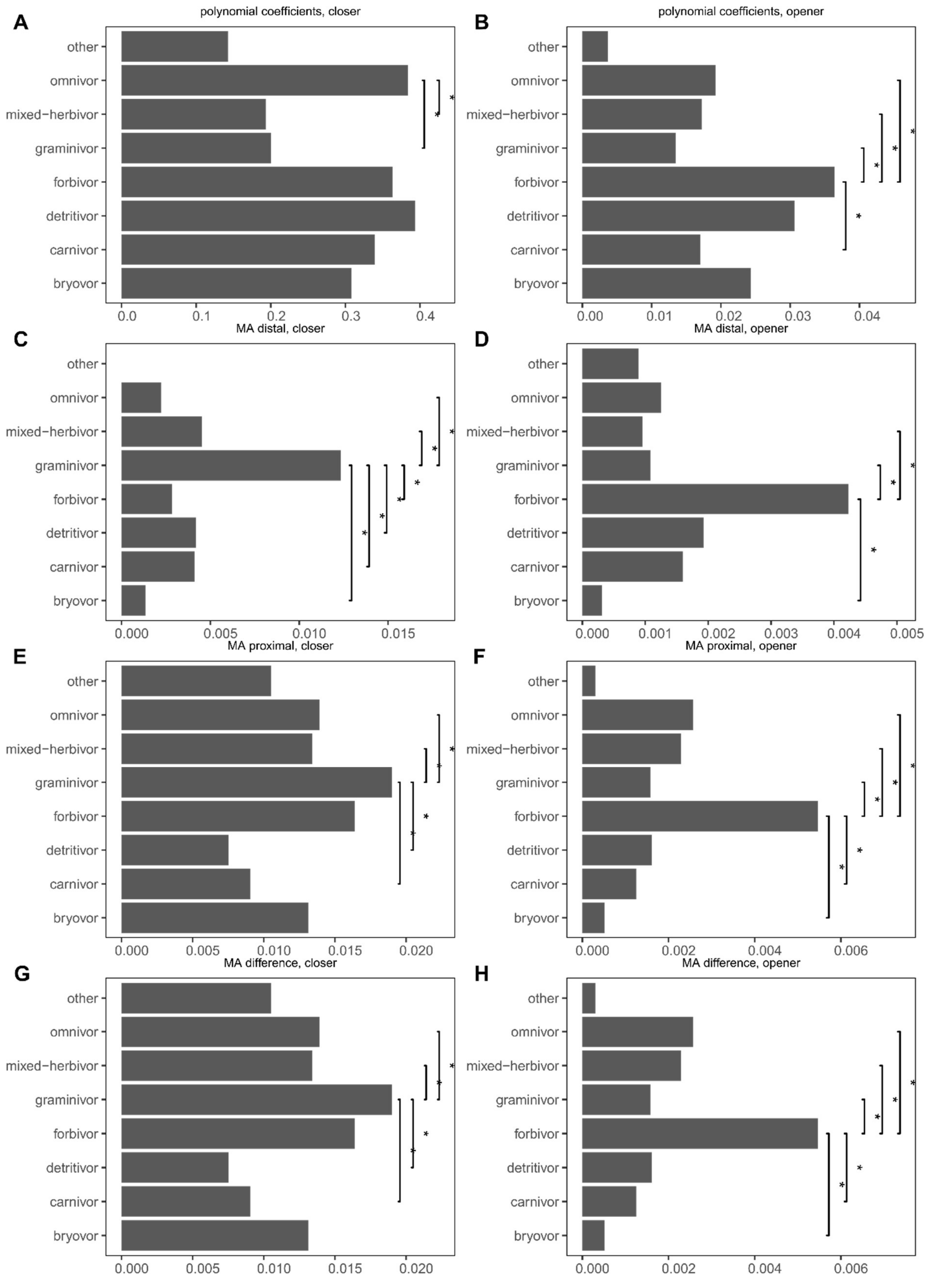

*S 9: Results of testing for disparity within diet superfamily groups for polynomial coefficients, distal MA, proximal, MA and MA difference on the closer and opener muscle Groups with significant difference of mean markes with *** = P<0.001; ** = P <0.01; P < 0.005*

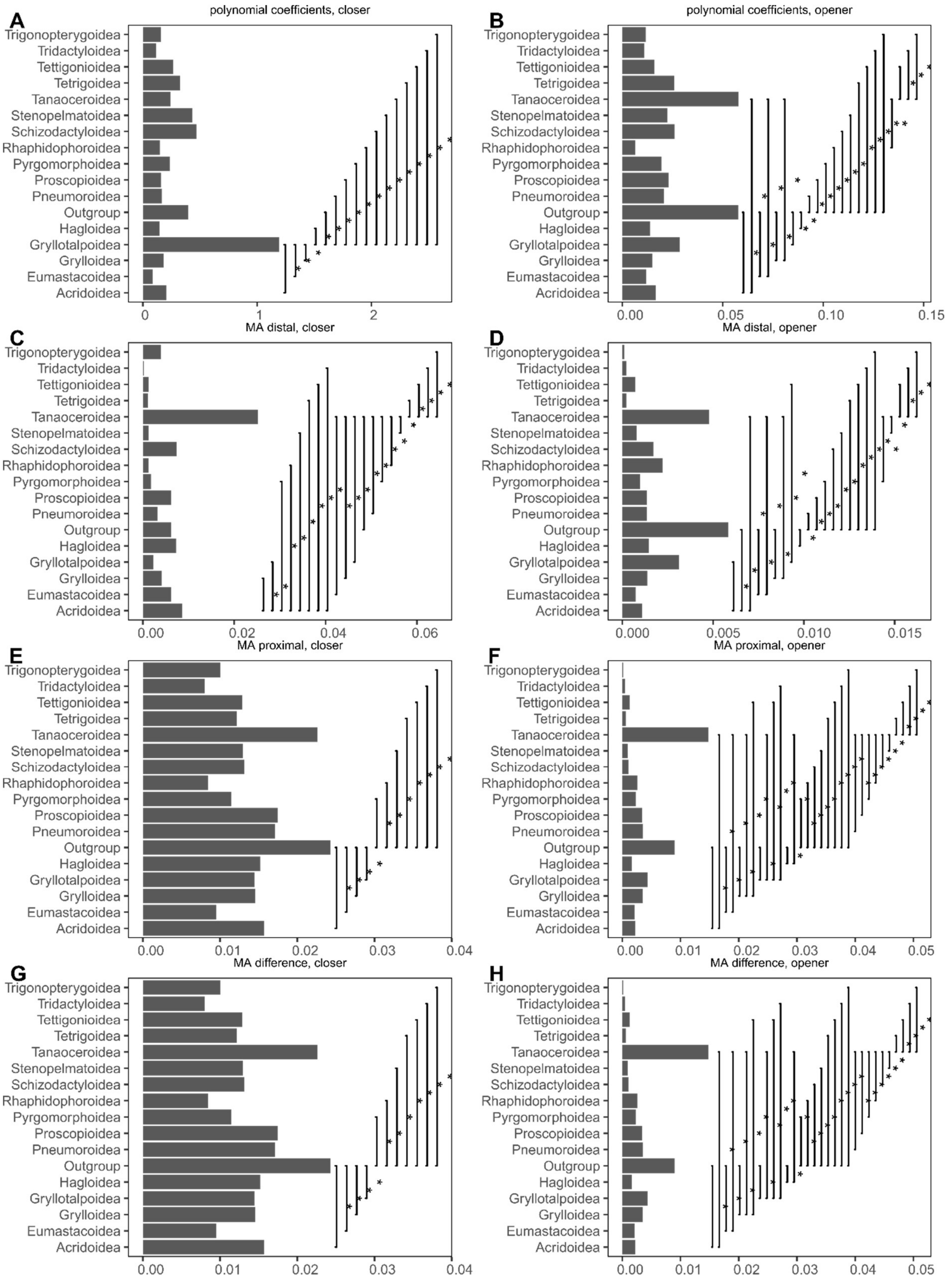
