## Supplement S10 for "Bite force transmission and mandible shape in grasshoppers, crickets, and allies is largely dependent on phylogeny, not diet"

*S 1: List of Species used in Analysis and acquired data*

| scientific Name | subset | order | suborder | superfamily | family | subfamily | genus | species | mandible length [mm] | GBIFspeciesKey | GBIFgenusKey | diet |
|---|---|---|---|---|---|---|---|---|---|---|---|---|
| Acrida bicolor | x | Orthoptera | Caelifera | Acridoidea | Acrididae | Acridinae | Acrida | bicolor | 2.94 | 1713654 | 1713619 | graminivor |
| Acrostegastes glaber | | Orthoptera | Caelifera | Acridoidea | Acrididae | Teratodinae | Acrostegastes | glaber | 1.6 | 1705304 | 1705302 | mixed-herbivor |
| Acrotylus insubricus | x | Orthoptera | Caelifera | Acridoidea | Acrididae | Oedipodinae | Acrotylus | insubricus | 2.11 | 5099079 | 1711256 | mixed-herbivor |
| Aeropedellus clavatus | | Orthoptera | Caelifera | Acridoidea | Acrididae | Gomphocerinae | Aeropedellus | clavatus | 3.12 | 1700119 | 1700116 | graminivor |
| Allotruxalis gracilis | | Orthoptera | Caelifera | Acridoidea | Acrididae | Acridinae | Allotruxalis | gracilis | 1.22 | 1703936 | 1703935 | graminivor |
| Amblytropidia australis | x | Orthoptera | Caelifera | Acridoidea | Acrididae | Gomphocerinae | Amblytropidia | australis | 2.87 | 1708033 | 1708013 | graminivor |
| Barombia tuberculosa | | Orthoptera | Caelifera | Acridoidea | Acrididae | Catantopinae | Barombia | tuberculosa | 2.4 | 5098964 | 1710400 | mixed-herbivor |
| Bruneria brunnea | x | Orthoptera | Caelifera | Acridoidea | Acrididae | Gomphocerinae | Bruneria | brunnea | 2.01 | 1704377 | 1700226 | graminivor |
| Cannula gracilis | x | Orthoptera | Caelifera | Acridoidea | Acrididae | Acridinae | Cannula | gracilis | 1.21 | 1705289 | 1705283 | mixed-herbivor |
| Carbonellacris grossa | x | Orthoptera | Caelifera | Acridoidea | Acrididae | Leptysminae | Carbonellacris | grossa | 1.72 | 1708948 | 1708947 | graminivor |
| Catantops melanostictus | x | Orthoptera | Caelifera | Acridoidea | Acrididae | Catantopinae | Catantops | melanostictus | 1.93 | 1710683 | 1710670 | mixed-herbivor |
| Catantops sp. | x | Orthoptera | Caelifera | Acridoidea | Acrididae | Catantopinae | Catantops | sp. | 2.07 | | 1710670 | mixed-herbivor |
| Chirista compta | | Orthoptera | Caelifera | Acridoidea | Acrididae | Acridinae | Chirista | compta | 0.6 | 1705077 | 1705076 | mixed-herbivor |
| Chloealtis abdominalis | x | Orthoptera | Caelifera | Acridoidea | Acrididae | Gomphocerinae | Chloealtis | abdominalis | 1.62 | 1701298 | 1701288 | graminivor |
| Chloropseustes brunneus | x | Orthoptera | Caelifera | Acridoidea | Acrididae | Leptysminae | Chloropseustes | brunneus | 0.66 | 1704588 | 1704574 | graminivor |
| Duroniella lucasii | x | Orthoptera | Caelifera | Acridoidea | Acrididae | Acridinae | Duroniella | lucasii | 1.25 | 1713168 | 1713166 | mixed-herbivor |
| Egnatioides sp. | x | Orthoptera | Caelifera | Acridoidea | Acrididae | Egnatiinae | Egnatioides | sp. | 1.38 | | 1701718 | mixed-herbivor |
| Eremogryllus hammadae | | Orthoptera | Caelifera | Acridoidea | Acrididae | Eremogryllinae | Eremogryllus | hammadae | 1 | 1701817 | 1701816 | mixed-herbivor |
| Euthystira brachyptera | x | Orthoptera | Caelifera | Acridoidea | Acrididae | Gomphocerinae | Euthystira | brachyptera | 1.59 | 1701343 | 1701338 | graminivor |
| Habrocnemis shanensis | | Orthoptera | Caelifera | Acridoidea | Acrididae | Habrocneminae | Habrocnemis | shanensis | 2.33 | 1706263 | 1706262 | mixed-herbivor |
| Kassongia maculifemur | x | Orthoptera | Caelifera | Acridoidea | Acrididae | Hemiacridinae | Kassongia | maculifemur | 1.53 | 1701825 | 1701820 | mixed-herbivor |
| Marellia remipes | x | Orthoptera | Caelifera | Acridoidea | Acrididae | Marelliinae | Marellia | remipes | 1.28 | 1711189 | 1711188 | mixed-herbivor |
| Mesopsis laticornis | x | Orthoptera | Caelifera | Acridoidea | Acrididae | Gomphocerinae | Mesopsis | laticornis | 2.13 | 1702997 | 1702987 | graminivor |
| Notostaurus anatolicus | x | Orthoptera | Caelifera | Acridoidea | Acrididae | Gomphocerinae | Notostaurus | anatolicus | 1.67 | 1704291 | 1704289 | graminivor |
| Odontomelus scalatus | | Orthoptera | Caelifera | Acridoidea | Acrididae | Acridinae | Odontomelus | scalatus | 1.91 | 1713213 | 1713212 | graminivor |
| Ommatolampis perspicillata | x | Orthoptera | Caelifera | Acridoidea | Acrididae | Ommatolampidinae | Ommatolampis | perspicillata | 1.74 | 1707767 | 1707765 | mixed-herbivor |
| Omocestus panteli | x | Orthoptera | Caelifera | Acridoidea | Acrididae | Gomphocerinae | Omocestus | panteli | 1.87 | 1710752 | 1710730 | graminivor |
| Parapropacris notatus | | Orthoptera | Caelifera | Acridoidea | Acrididae | Catantopinae | Parapropacris | notatus | 1.73 | 1711662 | 1711653 | mixed-herbivor |
| Parascopas sanguineus | x | Orthoptera | Caelifera | Acridoidea | Acrididae | Melanoplinae | Parascopas | sanguineus | 2.3 | 1702953 | 1702946 | mixed-herbivor |
| Paulinia acuminata | x | Orthoptera | Caelifera | Acridoidea | Acrididae | Pauliniinae | Paulinia | acuminata | 1.58 | 1710385 | 7760425 | forbivor |
| Phlibostroma quadrimaculatum | x | Orthoptera | Caelifera | Acridoidea | Acrididae | Gomphocerinae | Phlibostroma | quadrimaculatum | 2.29 | 1699988 | 1699987 | graminivor |
| Pnorisa squalus | x | Orthoptera | Caelifera | Acridoidea | Acrididae | Gomphocerinae | Pnorisa | squalus | 1.65 | 1704842 | 1704834 | graminivor |
| Podisma pedestris | x | Orthoptera | Caelifera | Acridoidea | Acrididae | Melanoplinae | Podisma | pedestris | 2.42 | 1704142 | 1697874 | graminivor |
| Pseudoxya diminuta | x | Orthoptera | Caelifera | Acridoidea | Acrididae | Oxyinae | Pseudoxya | diminuta | 1.26 | 1712766 | 1712765 | graminivor |
| Spathosternum prasiniferum | x | Orthoptera | Caelifera | Acridoidea | Acrididae | Spathosterninae | Spathosternum | prasiniferum | 1.04 | 5098610 | 1705766 | mixed-herbivor |
| Stauroderus scalaris | x | Orthoptera | Caelifera | Acridoidea | Acrididae | Gomphocerinae | Stauroderus | scalaris | 1.75 | 1703827 | 1703820 | graminivor |
| Tropidopola longicornis | x | Orthoptera | Caelifera | Acridoidea | Acrididae | Tropidopolinae | Tropidopola | longicornis | 1.34 | 1709410 | 1709408 | mixed-herbivor |
| Conophyma sogdianum | | Orthoptera | Caelifera | Acridoidea | Dericorythidae | Conophyminae | Conophyma | sogdianum | 1.58 | 1715649 | 1715582 | mixed-herbivor |
| Iranella eremiaphila | | Orthoptera | Caelifera | Acridoidea | Dericorythidae | Iranellinae | Iranella | eremiaphila | 1.51 | 5099761 | 10864554 | mixed-herbivor |

| | | | | | | | | | | | | |
|---|---|---|---|---|---|---|---|---|---|---|---|---|
| Iraniobia pavlovskii | | Orthoptera | Caelifera | Acridoidea | Dericorythidae | Iranellinae | Iraniobia | pavlovskii | 1.54 | 1715538 | 1715532 | mixed-herbivor |
| Batrachidacris rubridens | | Orthoptera | Caelifera | Acridoidea | Lathiceridae | | Batrachidacris | rubridens | 3.33 | 1715508 | 1715507 | mixed-herbivor |
| Lathicerus cimex | | Orthoptera | Caelifera | Acridoidea | Lathiceridae | | Lathicerus | cimex | 5.38 | 1715511 | 1715510 | mixed-herbivor |
| Afrotettix gariepensis | | Orthoptera | Caelifera | Acridoidea | Lentulidae | Shelforditinae | Afrotettix | gariepensis | 1.09 | 1715960 | 1715958 | mixed-herbivor |
| Betiscoides meridionalis | x | Orthoptera | Caelifera | Acridoidea | Lentulidae | Lentulinae | Betiscoides | meridionalis | 1.09 | 1715882 | 1715881 | mixed-herbivor |
| Shelfordites nanus | | Orthoptera | Caelifera | Acridoidea | Lentulidae | Shelforditinae | Shelfordites | nanus | 1.04 | 1715816 | 1715814 | mixed-herbivor |
| Eneremius punctifrons | x | Orthoptera | Caelifera | Acridoidea | Lithidiidae | | Eneremius | punctifrons | 0.8 | 8301826 | 1715488 | mixed-herbivor |
| Clarazella patagona | x | Orthoptera | Caelifera | Acridoidea | Ommexechidae | Ommexechinae | Clarazella | patagona | 1.3 | 1715420 | 1715409 | mixed-herbivor |
| Conometopus sulcaticollis | x | Orthoptera | Caelifera | Acridoidea | Ommexechidae | Aucacridinae | Conometopus | sulcaticollis | 1.89 | 5099746 | 1715442 | mixed-herbivor |
| Geloiomimus nasicus | | Orthoptera | Caelifera | Acridoidea | Pamphagidae | Echinotropinae | Geloiomimus | nasicus | 1.76 | 1697748 | 1697747 | mixed-herbivor |
| Glauia durieui | | Orthoptera | Caelifera | Acridoidea | Pamphagidae | Pamphaginae | Glauia | durieui | 2.3 | 1697536 | 1697534 | mixed-herbivor |
| Glyphanus obtusus | | Orthoptera | Caelifera | Acridoidea | Pamphagidae | Thrinchinae | Glyphanus | obtusus | 2.23 | 1697739 | 1697738 | mixed-herbivor |
| Haplotropis brunneriana | x | Orthoptera | Caelifera | Acridoidea | Pamphagidae | Thrinchinae | Haplotropis | brunneriana | 3.49 | 1697982 | 1697980 | forbivor |
| Ocnerosthenus kneuckeri | | Orthoptera | Caelifera | Acridoidea | Pamphagidae | Pamphaginae | Ocnerosthenus | kneuckeri | 0.4 | 1697523 | 1697520 | mixed-herbivor |
| Pamphagus sardeus | | Orthoptera | Caelifera | Acridoidea | Pamphagidae | Pamphaginae | Pamphagus | sardeus | 3.88 | 5098001 | 1698186 | forbivor |
| Prionotropis hystrix | x | Orthoptera | Caelifera | Acridoidea | Pamphagidae | Thrinchinae | Prionotropis | hystrix | 3.57 | 5098052 | 1698384 | forbivor |
| Thrincotropis caffra | | Orthoptera | Caelifera | Acridoidea | Pamphagidae | Echinotropinae | Thrincotropis | caffra | 1.5 | 1698626 | 1698624 | mixed-herbivor |
| Tmethis cisti | x | Orthoptera | Caelifera | Acridoidea | Pamphagidae | Thrinchinae | Tmethis | cisti | 3.45 | 1698059 | 1698051 | graminivor |
| Utubius syriacus | | Orthoptera | Caelifera | Acridoidea | Pamphagidae | Thrinchinae | Utubius | syriacus | 1.3 | 1698761 | 1698760 | mixed-herbivor |
| Charilaus carinatus | x | Orthoptera | Caelifera | Acridoidea | Pamphagodidae | | Charilaus | carinatus | 5.33 | 1715385 | 1715384 | mixed-herbivor |
| Chromacris icterus | x | Orthoptera | Caelifera | Acridoidea | Romaleidae | Romaleinae | Chromacris | icterus | 3.05 | 1714444 | 1714440 | mixed-herbivor |
| Helicopacris sp. | | Orthoptera | Caelifera | Acridoidea | Romaleidae | Bactrophorinae | Helicopacris | sp. | 2.17 | | 1715365 | mixed-herbivor |
| Ophthalmolampis sigillata | | Orthoptera | Caelifera | Acridoidea | Romaleidae | Bactrophorinae | Ophthalmolampis | sigillata | 2.45 | 1714784 | 1709564 | mixed-herbivor |
| Romalea microptera | x | Orthoptera | Caelifera | Acridoidea | Romaleidae | Romaleinae | Romalea | microptera | 3.44 | 1714599 | 1714598 | omnivor |
| Xyleus laevipes | x | Orthoptera | Caelifera | Acridoidea | Romaleidae | Romaleinae | Xyleus | laevipes | 3.09 | 1714433 | 1714350 | mixed-herbivor |
| Bufonacris terrestris | x | Orthoptera | Caelifera | Acridoidea | Tristiridae | Tristirinae | Bufonacris | terrestris | 3.45 | 1697467 | 1697463 | mixed-herbivor |
| Burrinia humbertiana | | Orthoptera | Caelifera | Eumastacoidea | Chorotypidae | Chorotypinae | Burrinia | humbertiana | 2.05 | 1696749 | 1696748 | mixed-herbivor |
| China mantispoides | | Orthoptera | Caelifera | Eumastacoidea | Chorotypidae | Chininae | China | mantispoides | 1.53 | 1696932 | 1696931 | mixed-herbivor |
| Chorotypus servillei | x | Orthoptera | Caelifera | Eumastacoidea | Chorotypidae | Chorotypinae | Chorotypus | servillei | 1.37 | 1696914 | 1696897 | mixed-herbivor |
| Erucius sp. | x | Orthoptera | Caelifera | Eumastacoidea | Chorotypidae | Eruciinae | Erucius | sp. | 1.16 | | 1696822 | mixed-herbivor |
| Orchetypus rugifrons | | Orthoptera | Caelifera | Eumastacoidea | Chorotypidae | Chorotypinae | Orchetypus | rugifrons | 1.63 | 1697012 | 1697009 | mixed-herbivor |
| Phyllochoreia unicolor | | Orthoptera | Caelifera | Eumastacoidea | Chorotypidae | Chorotypinae | Phyllochoreia | unicolor | 1.09 | 1696963 | 1696961 | mixed-herbivor |
| Prionacantha picta | | Orthoptera | Caelifera | Eumastacoidea | Chorotypidae | Prionacanthinae | Prionacantha | picta | 1.68 | 1696896 | 1696895 | mixed-herbivor |
| Pseudomnesicles rhodopeplus | | Orthoptera | Caelifera | Eumastacoidea | Chorotypidae | Mnesicleinae | Pseudomnesicles | rhodopeplus | 1.54 | 1696812 | 1696805 | mixed-herbivor |
| Uvarovia longipennis | | Orthoptera | Caelifera | Eumastacoidea | Chorotypidae | Mnesicleinae | Uvarovia | longipennis | 1.81 | 1696992 | 1696991 | mixed-herbivor |
| Heteromastax appendiculata | | Orthoptera | Caelifera | Eumastacoidea | Episactidae | Miraculinae | Heteromastax | appendiculata | 1.06 | 1695569 | 1695567 | mixed-herbivor |
| Lethus nicaraguae | x | Orthoptera | Caelifera | Eumastacoidea | Episactidae | Episactinae | Lethus | nicaraguae | 1.05 | 5097801 | 1695600 | mixed-herbivor |
| Malagassa tridens | | Orthoptera | Caelifera | Eumastacoidea | Episactidae | Miraculinae | Malagassa | tridens | 1.62 | 1695614 | 1695608 | mixed-herbivor |
| Pielomastax octavii | x | Orthoptera | Caelifera | Eumastacoidea | Episactidae | Episactinae | Pielomastax | octavii | 1.68 | 1695580 | 1695572 | mixed-herbivor |
| Seyrigella notabilis | | Orthoptera | Caelifera | Eumastacoidea | Episactidae | Miraculinae | Seyrigella | notabilis | 0.85 | 1695618 | 1695617 | mixed-herbivor |
| Teicophrys strigilecula | | Orthoptera | Caelifera | Eumastacoidea | Episactidae | Teicophryinae | Teicophrys | strigilecula | 1.1 | 1695595 | 1695585 | mixed-herbivor |
| Clinomastax ninae | | Orthoptera | Caelifera | Eumastacoidea | Eumastacidae | Gomphomastacinae | Clinomastax | ninae | 1.41 | 1697195 | 1697194 | mixed-herbivor |

| | | | | | | | | | | | | |
|---|---|---|---|---|---|---|---|---|---|---|---|---|
| Erythromastax kergarioui | | Orthoptera | Caelifera | Eumastacoidea | Eumastacidae | Eumastacinae | Erythromastax | kergarioui | 0.87 | 1697144 | 1697143 | mixed-herbivor |
| Eumastax sp. | | Orthoptera | Caelifera | Eumastacoidea | Eumastacidae | Eumastacinae | Eumastax | sp. | 0.94 | | 1697208 | mixed-herbivor |
| Gomphomastax antennata | | Orthoptera | Caelifera | Eumastacoidea | Eumastacidae | Gomphomastacinae | Gomphomastax | antennata | 1.22 | 1697287 | 1697285 | mixed-herbivor |
| Gomphomastax clavata | | Orthoptera | Caelifera | Eumastacoidea | Eumastacidae | Gomphomastacinae | Gomphomastax | clavata | 0.92 | 1697295 | 1697285 | mixed-herbivor |
| Morsea dumicola | | Orthoptera | Caelifera | Eumastacoidea | Eumastacidae | Morseinae | Morsea | dumicola | 1.13 | 1697266 | 1697265 | mixed-herbivor |
| Paramastax nigra | x | Orthoptera | Caelifera | Eumastacoidea | Eumastacidae | Paramastacinae | Paramastax | nigra | 1.28 | 1697095 | 1697087 | mixed-herbivor |
| Parepisactus carinatus | | Orthoptera | Caelifera | Eumastacoidea | Eumastacidae | Parepisactinae | Parepisactus | carinatus | 1.06 | 1697054 | 1697052 | mixed-herbivor |
| Pseudomastax laeta | | Orthoptera | Caelifera | Eumastacoidea | Eumastacidae | Pseudomastacinae | Pseudomastax | laeta | 1.88 | 1697162 | 1697147 | mixed-herbivor |
| Temnomastax tigris | | Orthoptera | Caelifera | Eumastacoidea | Eumastacidae | Temnomastacinae | Temnomastax | tigris | 1.04 | 1697334 | 1697332 | mixed-herbivor |
| Apteropeoedes wintreberti | | Orthoptera | Caelifera | Eumastacoidea | Euschmidtiidae | Pseudoschmidtiinae | Apteropeoedes | wintreberti | 1.95 | 1696448 | 1696440 | mixed-herbivor |
| Caenoschmidtia vittata | | Orthoptera | Caelifera | Eumastacoidea | Euschmidtiidae | Euschmidtiinae | Caenoschmidtia | vittata | 1.52 | 1696504 | 1696501 | mixed-herbivor |
| Chromomastax guttatifrons | | Orthoptera | Caelifera | Eumastacoidea | Euschmidtiidae | Pseudoschmidtiinae | Chromomastax | guttatifrons | 1.62 | 1696367 | 1696363 | mixed-herbivor |
| Euschmidtia cruciformis | | Orthoptera | Caelifera | Eumastacoidea | Euschmidtiidae | Euschmidtiinae | Euschmidtia | cruciformis | 1.47 | 1696622 | 1696618 | mixed-herbivor |
| Exophtalmomastax lucicola | | Orthoptera | Caelifera | Eumastacoidea | Euschmidtiidae | Pseudoschmidtiinae | Exophtalmomastax | lucicola | 1.81 | 1696495 | 1696492 | mixed-herbivor |
| Micromastax teteforti | | Orthoptera | Caelifera | Eumastacoidea | Euschmidtiidae | Pseudoschmidtiinae | Micromastax | teteforti | 1.33 | 1696724 | 1696719 | mixed-herbivor |
| Paraschmidtia burri | | Orthoptera | Caelifera | Eumastacoidea | Euschmidtiidae | Euschmidtiinae | Paraschmidtia | burri | 1.91 | 1696683 | 1696682 | mixed-herbivor |
| Paramastacides ramachendrai | | Orthoptera | Caelifera | Eumastacoidea | Mastacideidae | Mastacideinae | Paramastacides | ramachendrai | 1.31 | 1695631 | 1695628 | mixed-herbivor |
| Biroella gracilis | | Orthoptera | Caelifera | Eumastacoidea | Morabidae | Biroellinae | Biroella | gracilis | 1.14 | 5097763 | 1695425 | mixed-herbivor |
| Heide amiculi | | Orthoptera | Caelifera | Eumastacoidea | Morabidae | Morabinae | Heide | amiculi | 0.72 | 1695416 | 1695411 | mixed-herbivor |
| Acanthothericles rubriventris | | Orthoptera | Caelifera | Eumastacoidea | Thericleidae | Chromothericleinae | Acanthothericles | rubriventris | 0.9 | 1695738 | 1695737 | mixed-herbivor |
| Afromastax zebra | | Orthoptera | Caelifera | Eumastacoidea | Thericleidae | Afromastacinae | Afromastax | zebra | 0.95 | 1695956 | 1695949 | mixed-herbivor |
| Bunkeya congoensis | | Orthoptera | Caelifera | Eumastacoidea | Thericleidae | Thericleinae | Bunkeya | congoensis | 1.88 | 1695674 | 1695673 | mixed-herbivor |
| Chromothericles keniensis | | Orthoptera | Caelifera | Eumastacoidea | Thericleidae | Chromothericleinae | Chromothericles | keniensis | 0.87 | 1695902 | 1695897 | forbivor |
| Phaulotypus socotranus | | Orthoptera | Caelifera | Eumastacoidea | Thericleidae | Plagiotriptinae | Phaulotypus | socotranus | 1.13 | 1695748 | 1695741 | mixed-herbivor |
| Thamithericles birunga | | Orthoptera | Caelifera | Eumastacoidea | Thericleidae | Afromastacinae | Thamithericles | birunga | 1.29 | 1695915 | 1695914 | mixed-herbivor |
| Thericles obtusifrons | x | Orthoptera | Caelifera | Eumastacoidea | Thericleidae | Thericleinae | Thericles | obtusifrons | 1.77 | 1695732 | 1695726 | mixed-herbivor |
| Thericlesiella meridionalis | | Orthoptera | Caelifera | Eumastacoidea | Thericleidae | Afromastacinae | Thericlesiella | meridionalis | 1.23 | 1695670 | 1695669 | mixed-herbivor |
| Physophorina livingstonii | x | Orthoptera | Caelifera | Pneumoroidea | Pneumoridae | | Physophorina | livingstonii | 3.32 | 1715981 | 1715980 | omnivor |
| Pneumora inanis | x | Orthoptera | Caelifera | Pneumoroidea | Pneumoridae | | Pneumora | inanis | 5.43 | 1716006 | 7775060 | mixed-herbivor |
| Hybusa occidentalis | x | Orthoptera | Caelifera | Proscopioidea | Proscopiidae | Hybusinae | Hybusa | occidentalis | 1.6 | 1696080 | 1696075 | mixed-herbivor |
| Proscopia sp. | x | Orthoptera | Caelifera | Proscopioidea | Proscopiidae | Proscopiinae | Proscopia | sp. | 4.03 | | 1696322 | mixed-herbivor |
| Caprorhinus zolotarevskyi | x | Orthoptera | Caelifera | Pyrgomorphoidea | Pyrgomorphidae | Orthacridinae | Caprorhinus | zolotarevskyi | 2 | 1727853 | 1727851 | forbivor |
| Gymnohippus marmoratus | x | Orthoptera | Caelifera | Pyrgomorphoidea | Pyrgomorphidae | Orthacridinae | Gymnohippus | marmoratus | 1.49 | 1728777 | 1728776 | mixed-herbivor |
| Phymateus sp. | x | Orthoptera | Caelifera | Pyrgomorphoidea | Pyrgomorphidae | Pyrgomorphinae | Phymateus | sp. | 3.21 | | 1728291 | mixed-herbivor |
| Phyteumas purpurascens | x | Orthoptera | Caelifera | Pyrgomorphoidea | Pyrgomorphidae | Pyrgomorphinae | Phyteumas | purpurascens | 2.71 | 1728257 | 1728256 | mixed-herbivor |
| Rutidoderes squarrosus | x | Orthoptera | Caelifera | Pyrgomorphoidea | Pyrgomorphidae | Pyrgomorphinae | Rutidoderes | squarrosus | 3.44 | 1728152 | 1728150 | mixed-herbivor |
| Taphronota calliparea | x | Orthoptera | Caelifera | Pyrgomorphoidea | Pyrgomorphidae | Pyrgomorphinae | Taphronota | calliparea | 2.52 | 1728712 | 1728700 | mixed-herbivor |
| Zonocerus elegans | x | Orthoptera | Caelifera | Pyrgomorphoidea | Pyrgomorphidae | Pyrgomorphinae | Zonocerus | elegans | 2.68 | 1727809 | 1700561 | forbivor |
| Tanaocerus koebelei | x | Orthoptera | Caelifera | Tanaoceroidea | Tanaoceridae | Tanaocerinae | Tanaocerus | koebelei | 0.82 | 1715976 | 1715975 | mixed-herbivor |
| Boczkitettix borneensis | | Orthoptera | Caelifera | Tetrigoidea | Tetrigidae | Cladonotinae | Boczkitettix | borneensis | 1.1 | 7781695 | 8231659 | bryovor |
| Bufonides uvarovi | | Orthoptera | Caelifera | Tetrigoidea | Tetrigidae | Batrachideinae | Bufonides | uvarovi | 0.78 | 1679700 | 1679698 | bryovor |
| Cleostratus monocerus | | Orthoptera | Caelifera | Tetrigoidea | Tetrigidae | Metrodorinae | Cleostratus | monocerus | 0.87 | 1680382 | 1680379 | bryovor |

| | | | | | | | | | | | | |
|---|---|---|---|---|---|---|---|---|---|---|---|---|
| Dinotettix sp. | x | Orthoptera | Caelifera | Tetrigoidea | Tetrigidae | Tetriginae | Dinotettix | sp. | 0.63 | | 1682222 | mixed-herbivor |
| Discotettix belzebuth | | Orthoptera | Caelifera | Tetrigoidea | Tetrigidae | Scelimeninae | Discotettix | belzebuth | 1.5 | 1679555 | 1679552 | bryovor |
| Eomorphopus antennatus | | Orthoptera | Caelifera | Tetrigoidea | Tetrigidae | Metrodorinae | Eomorphopus | antennatus | 1.17 | 1680193 | 1680191 | bryovor |
| Ergatettix dorsifera | | Orthoptera | Caelifera | Tetrigoidea | Tetrigidae | Tetriginae | Ergatettix | dorsifera | 0.93 | 4713206 | 1682058 | bryovor |
| Eutettigidea lineata | | Orthoptera | Caelifera | Tetrigoidea | Tetrigidae | Batrachideinae | Eutettigidea | lineata | 0.85 | 1679704 | 1679703 | bryovor |
| Holocerus lucifer | | Orthoptera | Caelifera | Tetrigoidea | Tetrigidae | Metrodorinae | Holocerus | lucifer | 1.17 | 1682032 | 1682030 | bryovor |
| Hymenotes triangularis | | Orthoptera | Caelifera | Tetrigoidea | Tetrigidae | Cladonotinae | Hymenotes | triangularis | 1.41 | 1679901 | 1679899 | bryovor |
| Lamellitettigodes sagittatus | x | Orthoptera | Caelifera | Tetrigoidea | Tetrigidae | Tetriginae | Lamellitettigodes | sagittatus | 0.47 | 9961019 | 1682211 | bryovor |
| Leptacrydium sp. | x | Orthoptera | Caelifera | Tetrigoidea | Tetrigidae | Tetriginae | Leptacrydium | sp. | 0.69 | | 1680996 | bryovor |
| Loxilobus leveri | | Orthoptera | Caelifera | Tetrigoidea | Tetrigidae | | Loxilobus | leveri | 0.7 | 1680031 | 1680014 | bryovor |
| Macquillania pacifica | | Orthoptera | Caelifera | Tetrigoidea | Tetrigidae | Tetriginae | Macquillania | pacifica | 0.63 | 1682203 | 1682201 | mixed-herbivor |
| Micronotus quadriundulatus | | Orthoptera | Caelifera | Tetrigoidea | Tetrigidae | Tetriginae | Micronotus | quadriundulatus | 0.64 | 1680692 | 1680685 | bryovor |
| Neotettix femoratus | | Orthoptera | Caelifera | Tetrigoidea | Tetrigidae | Tetriginae | Neotettix | femoratus | 0.85 | 1680911 | 1680907 | bryovor |
| Paratettix amplus | | Orthoptera | Caelifera | Tetrigoidea | Tetrigidae | Tetriginae | Paratettix | amplus | 0.78 | 1680591 | 1680419 | bryovor |
| Platygavialidium dentifer | | Orthoptera | Caelifera | Tetrigoidea | Tetrigidae | Scelimeninae | Platygavialidium | dentifer | 1.31 | 1681598 | 1681592 | bryovor |
| Pseudoparatettix difficilis | | Orthoptera | Caelifera | Tetrigoidea | Tetrigidae | Metrodorinae | Pseudoparatettix | difficilis | 0.84 | 1680319 | 1680311 | bryovor |
| Pterotettix bigibbosus | | Orthoptera | Caelifera | Tetrigoidea | Tetrigidae | Metrodorinae | Pterotettix | bigibbosus | 1.38 | 1680681 | 1680669 | bryovor |
| Rostella processus | | Orthoptera | Caelifera | Tetrigoidea | Tetrigidae | Metrodorinae | Rostella | processus | 0.56 | 8828769 | 1679959 | bryovor |
| Tripetalocera ferruginea | | Orthoptera | Caelifera | Tetrigoidea | Tetrigidae | Tripetalocerinae | Tripetalocera | ferruginea | 1.03 | 1680849 | 1680847 | bryovor |
| Trypophyllum glabrifrons | x | Orthoptera | Caelifera | Tetrigoidea | Tetrigidae | Cladonotinae | Trypophyllum | glabrifrons | 0.76 | 1679894 | 1679893 | bryovor |
| Xistrella dromadaria | | Orthoptera | Caelifera | Tetrigoidea | Tetrigidae | Metrodorinae | Xistrella | dromadaria | 0.56 | 1681918 | 1681912 | bryovor |
| Mirhipipteryx columbiana | x | Orthoptera | Caelifera | Tridactyloidea | Ripipterygidae | Ripipteryginae | Mirhipipteryx | columbiana | 0.41 | 1724579 | 1724573 | omnivor |
| Ripipteryx atra | x | Orthoptera | Caelifera | Tridactyloidea | Ripipterygidae | Ripipteryginae | Ripipteryx | atra | 0.57 | 1724650 | 1724623 | omnivor |
| Afrotridactylus meridianus | x | Orthoptera | Caelifera | Tridactyloidea | Tridactylidae | Tridactylinae | Afrotridactylus | meridianus | 0.7 | 1724961 | 1724956 | omnivor |
| Asiotridactylus fasciatus | | Orthoptera | Caelifera | Tridactyloidea | Tridactylidae | Tridactylinae | Asiotridactylus | fasciatus | 0.41 | 1724857 | 1724856 | omnivor |
| Bruntridactylus brunneri | | Orthoptera | Caelifera | Tridactyloidea | Tridactylidae | Dentridactylinae | Bruntridactylus | brunneri | 0.86 | 1724884 | 1724877 | omnivor |
| Bruntridactylus irremipes | | Orthoptera | Caelifera | Tridactyloidea | Tridactylidae | Dentridactylinae | Bruntridactylus | irremipes | 0.78 | 1724889 | 1724877 | omnivor |
| Neotridactylus apicalis | x | Orthoptera | Caelifera | Tridactyloidea | Tridactylidae | Tridactylinae | Neotridactylus | apicalis | 0.74 | 1724825 | 1724819 | omnivor |
| Tridactylus thoracicus | x | Orthoptera | Caelifera | Tridactyloidea | Tridactylidae | Tridactylinae | Tridactylus | thoracicus | 0.83 | 5100800 | 1724686 | omnivor |
| Systella rafflesii | x | Orthoptera | Caelifera | Trigonopterygoidea | Trigonopterygidae | Trigonopteryginae | Systella | rafflesii | 2 | 1727358 | 1727347 | mixed-herbivor |
| Trigonopteryx hopei | x | Orthoptera | Caelifera | Trigonopterygoidea | Trigonopterygidae | Trigonopteryginae | Trigonopteryx | hopei | 1.97 | 1727375 | 1727372 | mixed-herbivor |
| Xyronotus aztecus | x | Orthoptera | Caelifera | Trigonopterygoidea | Xyronotidae | Xyronotinae | Xyronotus | aztecus | 0.9 | 1727380 | 1727379 | mixed-herbivor |
| Amblyrhethus brevipes | | Orthoptera | Ensifera | Grylloidea | Gryllidae | Oecanthinae | Amblyrhethus | brevipes | 1.49 | 1719983 | 1719976 | omnivor |
| Anurogryllus muticus | | Orthoptera | Ensifera | Grylloidea | Gryllidae | Gryllinae | Anurogryllus | muticus | 2.47 | 1717586 | 1717574 | omnivor |
| Aphemogryllus gracillis | | Orthoptera | Ensifera | Grylloidea | Gryllidae | Pentacentrinae | Aphemogryllus | gracillis | 0.79 | 1721654 | 1721653 | omnivor |
| Burrianus pachyceros | | Orthoptera | Ensifera | Grylloidea | Gryllidae | Euscyrtinae | Burrianus | pachyceros | 0.66 | 1716893 | 1716892 | omnivor |
| Eugryllodes littoreus | | Orthoptera | Ensifera | Grylloidea | Gryllidae | Gryllomorphinae | Eugryllodes | littoreus | 0.82 | 1716615 | 1716589 | omnivor |
| Eugryllodes pipiens | | Orthoptera | Ensifera | Grylloidea | Gryllidae | Gryllomorphinae | Eugryllodes | pipiens | 1.8 | 1716590 | 1716589 | omnivor |
| Euscyrtus bivittatus | | Orthoptera | Ensifera | Grylloidea | Gryllidae | Euscyrtinae | Euscyrtus | bivittatus | 0.54 | 1720515 | 1720499 | omnivor |
| Gryllomimus chopardi | | Orthoptera | Ensifera | Grylloidea | Gryllidae | Gryllomiminae | Gryllomimus | chopardi | 0.83 | 6543691 | 1723177 | omnivor |
| Gryllopsis sp. | | Orthoptera | Ensifera | Grylloidea | Gryllidae | Gryllinae | Gryllopsis | sp. | 1.94 | | 1716957 | omnivor |
| Gryllus bimaculatus | x | Orthoptera | Ensifera | Grylloidea | Gryllidae | Gryllinae | Gryllus | bimaculatus | 3.56 | 1713034 | 1404683 | omnivor |

| | | | | | | | | | | | | |
|---|---|---|---|---|---|---|---|---|---|---|---|---|
| Gryllus maunus | x | Orthoptera | Ensifera | Grylloidea | Gryllidae | Gryllinae | Gryllus | maunus | 1.04 | 1716785 | 1404683 | omnivor |
| Hapithus agitator | | Orthoptera | Ensifera | Grylloidea | Gryllidae | Hapithinae | Hapithus | agitator | 0.96 | 1717340 | 1717319 | omnivor |
| Hapithus similis | | Orthoptera | Ensifera | Grylloidea | Gryllidae | Hapithinae | Hapithus | similis | 1.29 | 9469760 | 1717319 | omnivor |
| Itara sonabilis | | Orthoptera | Ensifera | Grylloidea | Gryllidae | Itarinae | Itara | sonabilis | 1.52 | 1719101 | 1719050 | omnivor |
| Landreva clara | | Orthoptera | Ensifera | Grylloidea | Gryllidae | Landrevinae | Landreva | clara | 1.96 | 1716366 | 1716361 | omnivor |
| Leptogryllus similis | | Orthoptera | Ensifera | Grylloidea | Gryllidae | Oecanthinae | Leptogryllus | similis | 1 | 1717539 | 1717512 | omnivor |
| Loxoblemmus chopardi | x | Orthoptera | Ensifera | Grylloidea | Gryllidae | Gryllinae | Loxoblemmus | chopardi | 3.99 | 1721403 | 1721341 | omnivor |
| Melanogryllus desertus | | Orthoptera | Ensifera | Grylloidea | Gryllidae | Gryllinae | Melanogryllus | desertus | 2.26 | 1718616 | 1718612 | omnivor |
| Mitius blennus | | Orthoptera | Ensifera | Grylloidea | Gryllidae | Gryllinae | Mitius | blennus | 1.5 | 1721038 | 1721029 | omnivor |
| Modicogryllus confirmatus | x | Orthoptera | Ensifera | Grylloidea | Gryllidae | Gryllinae | Modicogryllus | confirmatus | 1.72 | 1722166 | 1722085 | omnivor |
| Myara sordida | | Orthoptera | Ensifera | Grylloidea | Gryllidae | Eneopterinae | Myara | sordida | 1.11 | 1722602 | 1722600 | omnivor |
| Nisitrus vittatus | | Orthoptera | Ensifera | Grylloidea | Gryllidae | Eneopterinae | Nisitrus | vittatus | 1.36 | 1716539 | 1716531 | other |
| Oecanthus pellucens | x | Orthoptera | Ensifera | Grylloidea | Gryllidae | Oecanthinae | Oecanthus | pellucens | 0.7 | 1720967 | 1699510 | omnivor |
| Parapentacentrus fuscus | | Orthoptera | Ensifera | Grylloidea | Gryllidae | Itarinae | Parapentacentrus | fuscus | 1.29 | 1718487 | 1718483 | omnivor |
| Pentacentrodes tenellus | | Orthoptera | Ensifera | Grylloidea | Gryllidae | Pentacentrinae | Pentacentrodes | tenellus | 0.55 | 1721651 | 1721650 | omnivor |
| Phonarellus lucens | | Orthoptera | Ensifera | Grylloidea | Gryllidae | Gryllinae | Phonarellus | lucens | 2.63 | 1723215 | 1723214 | omnivor |
| Plebeiogryllus guttiventris | x | Orthoptera | Ensifera | Grylloidea | Gryllidae | Gryllinae | Plebeiogryllus | guttiventris | 2.76 | 1723786 | 1723775 | omnivor |
| Rupilius nigrosignatus | | Orthoptera | Ensifera | Grylloidea | Gryllidae | Podoscirtinae | Rupilius | nigrosignatus | 1.46 | 1721982 | 1721981 | mixed-herbivor |
| Scapsipedus marginatus | | Orthoptera | Ensifera | Grylloidea | Gryllidae | Gryllinae | Scapsipedus | marginatus | 2.58 | 1723601 | 1723591 | omnivor |
| Sciobia caliendra | x | Orthoptera | Ensifera | Grylloidea | Gryllidae | Gryllinae | Sciobia | caliendra | 2.82 | 5100452 | 1722412 | omnivor |
| Sciobia lusitanica | x | Orthoptera | Ensifera | Grylloidea | Gryllidae | Gryllinae | Sciobia | lusitanica | 2.45 | 1722514 | 1722412 | mixed-herbivor |
| Sclerogryllus punctatus | | Orthoptera | Ensifera | Grylloidea | Gryllidae | Sclerogryllinae | Sclerogryllus | punctatus | 0.89 | 1723100 | 1723092 | omnivor |
| Teleogryllus occipitalis | x | Orthoptera | Ensifera | Grylloidea | Gryllidae | Gryllinae | Teleogryllus | occipitalis | 3.02 | 1723510 | 1723420 | omnivor |
| Trelleora sp. | | Orthoptera | Ensifera | Grylloidea | Gryllidae | Podoscirtinae | Trelleora | sp. | 1.48 | | 1719329 | mixed-herbivor |
| Truljalia citri | | Orthoptera | Ensifera | Grylloidea | Gryllidae | Podoscirtinae | Truljalia | citri | 1.16 | 1723286 | 1723274 | mixed-herbivor |
| Tumpalia ruficeps | | Orthoptera | Ensifera | Grylloidea | Gryllidae | Gryllinae | Tumpalia | ruficeps | 2.55 | 1721941 | 1721939 | omnivor |
| Xenogryllus eneopteroides | | Orthoptera | Ensifera | Grylloidea | Gryllidae | Eneopterinae | Xenogryllus | eneopteroides | 1.66 | 1720900 | 1720891 | omnivor |
| Zvenella yunnana | | Orthoptera | Ensifera | Grylloidea | Gryllidae | Podoscirtinae | Zvenella | yunnana | 1.2 | 1720276 | 1720259 | mixed-herbivor |
| Arachnocephalus vestitus | | Orthoptera | Ensifera | Grylloidea | Mogoplistidae | Mogoplistinae | Arachnocephalus | vestitus | 0.44 | 1724050 | 1724034 | omnivor |
| Cycloptiloides canariensis | | Orthoptera | Ensifera | Grylloidea | Mogoplistidae | Mogoplistinae | Cycloptiloides | canariensis | 0.5 | 1724266 | 1724256 | omnivor |
| Micrornebius incertus | | Orthoptera | Ensifera | Grylloidea | Mogoplistidae | Mogoplistinae | Micrornebius | incertus | 0.18 | 1724222 | 1724219 | omnivor |
| Ornebius marginatus | | Orthoptera | Ensifera | Grylloidea | Mogoplistidae | Mogoplistinae | Ornebius | marginatus | 0.23 | 1724405 | 1722339 | omnivor |
| Pseudomogoplistes squamiger | | Orthoptera | Ensifera | Grylloidea | Mogoplistidae | Mogoplistinae | Pseudomogoplistes | squamiger | 0.56 | 1724091 | 1724089 | omnivor |
| Aclodes chamocoru | | Orthoptera | Ensifera | Grylloidea | Phalangopsidae | Paragryllinae | Aclodes | chamocoru | 1.22 | 1719858 | 1719833 | omnivor |
| Amphiacusta annulipes | | Orthoptera | Ensifera | Grylloidea | Phalangopsidae | Luzarinae | Amphiacusta | annulipes | 1.6 | 1718934 | 1718857 | forbivor |
| Endacusta irrorata | | Orthoptera | Ensifera | Grylloidea | Phalangopsidae | Phalangopsinae | Endacusta | irrorata | 1.65 | 1720069 | 1720053 | omnivor |
| Endecous arachnopsis | | Orthoptera | Ensifera | Grylloidea | Phalangopsidae | Phalangopsinae | Endecous | arachnopsis | 1.2 | 1716645 | 1716631 | omnivor |
| Homoeogryllus venosus | | Orthoptera | Ensifera | Grylloidea | Phalangopsidae | Cachoplistinae | Homoeogryllus | venosus | 1.54 | 1720181 | 1720180 | omnivor |
| Homoeogryllus xanthographus | | Orthoptera | Ensifera | Grylloidea | Phalangopsidae | Cachoplistinae | Homoeogryllus | xanthographus | 1.57 | 1720194 | 1720180 | omnivor |
| Meloimorpha japonica | | Orthoptera | Ensifera | Grylloidea | Phalangopsidae | Cachoplistinae | Meloimorpha | japonica | 1.04 | 1719343 | 1719337 | omnivor |
| Paragryllus sp. | | Orthoptera | Ensifera | Grylloidea | Phalangopsidae | Paragryllinae | Paragryllus | sp. | 1.32 | | 1716231 | omnivor |
| Phaeophilacris bredoides | x | Orthoptera | Ensifera | Grylloidea | Phalangopsidae | Phalangopsinae | Phaeophilacris | bredoides | 1.97 | 1721748 | 1721671 | omnivor |

| | | | | | | | | | | | | |
|---|---|---|---|---|---|---|---|---|---|---|---|---|
| Phaeophilacris cavicola | x | Orthoptera | Ensifera | Grylloidea | Phalangopsidae | Phalangopsinae | Phaeophilacris | cavicola | 3.82 | 1721752 | 1721671 | omnivor |
| Phaeophilacris phalangium | | Orthoptera | Ensifera | Grylloidea | Phalangopsidae | Phalangopsinae | Phaeophilacris | phalangium | 0.45 | 1721734 | 1721671 | omnivor |
| Phaloria solomonica | | Orthoptera | Ensifera | Grylloidea | Phalangopsidae | Phaloriinae | Phaloria | solomonica | 1.12 | 1718089 | 1718038 | omnivor |
| Schizotrypus planus | | Orthoptera | Ensifera | Grylloidea | Phalangopsidae | Phaloriinae | Schizotrypus | planus | 1.2 | 1719022 | 1719021 | omnivor |
| Trellius buqueti | | Orthoptera | Ensifera | Grylloidea | Phalangopsidae | Phaloriinae | Trellius | buqueti | 1.33 | 1720611 | 1720593 | omnivor |
| Argizala brasilensis | | Orthoptera | Ensifera | Grylloidea | Trigonidiidae | Nemobiinae | Argizala | brasilensis | 0.61 | 1718262 | 1718258 | omnivor |
| Cranistus colliurides | | Orthoptera | Ensifera | Grylloidea | Trigonidiidae | Trigonidiinae | Cranistus | colliurides | 0.84 | 1716288 | 1716281 | mixed-herbivor |
| Dianemobius fascipes | x | Orthoptera | Ensifera | Grylloidea | Trigonidiidae | Nemobiinae | Dianemobius | fascipes | 0.32 | 1722867 | 1722854 | omnivor |
| Phylloscyrtus cicindeloides | | Orthoptera | Ensifera | Grylloidea | Trigonidiidae | Trigonidiinae | Phylloscyrtus | cicindeloides | 0.77 | 1720449 | 1720437 | mixed-herbivor |
| Pteronemobius heydenii | | Orthoptera | Ensifera | Grylloidea | Trigonidiidae | Nemobiinae | Pteronemobius | heydenii | 0.38 | 1717828 | 1717741 | omnivor |
| Tahitinemobius tigrinus | | Orthoptera | Ensifera | Grylloidea | Trigonidiidae | Nemobiinae | Tahitinemobius | tigrinus | 0.68 | 1719478 | 1719477 | omnivor |
| Trigonidium grande | x | Orthoptera | Ensifera | Grylloidea | Trigonidiidae | Trigonidiinae | Zudella | grande | 0.76 | 1721085 | 10800896 | mixed-herbivor |
| Pteroplistes sp. | | Orthoptera | Ensifera | Grylloidea | | Pteroplistinae | Pteroplistes | sp. | 1.15 | | 1722368 | omnivor |
| Gryllotalpa africana | x | Orthoptera | Ensifera | Gryllotalpoidea | Gryllotalpidae | Gryllotalpinae | Gryllotalpa | africana | 2.54 | 1716185 | 1716116 | omnivor |
| Gryllotalpa hirsuta | x | Orthoptera | Ensifera | Gryllotalpoidea | Gryllotalpidae | Gryllotalpinae | Gryllotalpa | hirsuta | 4.35 | 1716159 | 1716116 | omnivor |
| Gryllotalpella macilenta | x | Orthoptera | Ensifera | Gryllotalpoidea | Gryllotalpidae | Gryllotalpinae | Gryllotalpella | macilenta | 1.23 | 1716217 | 1716214 | omnivor |
| Neocurtilla hexadactyla | x | Orthoptera | Ensifera | Gryllotalpoidea | Gryllotalpidae | Gryllotalpinae | Neocurtilla | hexadactyla | 2.33 | 1716066 | 1716061 | omnivor |
| Scapteriscus oxydactylus | x | Orthoptera | Ensifera | Gryllotalpoidea | Gryllotalpidae | Scapteriscinae | Scapteriscus | oxydactylus | 3.76 | 1716106 | 1716084 | omnivor |
| Myrmecophilus acervorum | x | Orthoptera | Ensifera | Gryllotalpoidea | Myrmecophilidae | Myrmecophilinae | Myrmecophilus | acervorum | 0.29 | 5100649 | 1724436 | detritivor |
| Myrmecophilus hirticaudus | x | Orthoptera | Ensifera | Gryllotalpoidea | Myrmecophilidae | Myrmecophilinae | Myrmecophilus | hirticaudus | 0.35 | 5100639 | 1724436 | detritivor |
| Myrmecophilus ochraceus | x | Orthoptera | Ensifera | Gryllotalpoidea | Myrmecophilidae | Myrmecophilinae | Myrmecophilus | ochraceus | 0.28 | 5100666 | 1724436 | detritivor |
| Cyphoderris monstrosa | x | Orthoptera | Ensifera | Hagloidea | Prophalangopsidae | Cyphoderrinae | Cyphoderris | monstrosa | 4.01 | 1682545 | 1682543 | forbivor |
| Cyphoderris strepitans | x | Orthoptera | Ensifera | Hagloidea | Prophalangopsidae | Cyphoderrinae | Cyphoderris | strepitans | 4.83 | 1682549 | 1682543 | omnivor |
| Ceuthophilus pallidipes | x | Orthoptera | Ensifera | Rhaphidophoroidea | Rhaphidophoridae | Ceuthophilinae | Ceuthophilus | pallidipes | 1.32 | 1729692 | 1725135 | omnivor |
| Dolichopoda palpata | | Orthoptera | Ensifera | Rhaphidophoroidea | Rhaphidophoridae | Dolichopodainae | Dolichopoda | palpata | 2.42 | 1729270 | 1729247 | omnivor |
| Gammarotettix bilobatus | | Orthoptera | Ensifera | Rhaphidophoroidea | Rhaphidophoridae | Gammarotettiginae | Gammarotettix | bilobatus | 1.36 | 1729745 | 1729743 | omnivor |
| Pachyrhamma waitomoensis | x | Orthoptera | Ensifera | Rhaphidophoroidea | Rhaphidophoridae | Macropathinae | Pachyrhamma | waitomoensis | 3.41 | 1728911 | 1728884 | omnivor |
| Stonychophora papua | | Orthoptera | Ensifera | Rhaphidophoroidea | Rhaphidophoridae | Rhaphidophorinae | Stonychophora | papua | 2.2 | 1729232 | 1729203 | omnivor |
| Tachycines asynamorus | x | Orthoptera | Ensifera | Rhaphidophoroidea | Rhaphidophoridae | Aemodogryllinae | Tachycines | asynamorus | 2.24 | 1728965 | 1729113 | omnivor |
| Troglophilus cavicola | x | Orthoptera | Ensifera | Rhaphidophoroidea | Rhaphidophoridae | Troglophilinae | Troglophilus | cavicola | 1.99 | 1729803 | 1729796 | omnivor |
| Troglophilus neglectus | x | Orthoptera | Ensifera | Rhaphidophoroidea | Rhaphidophoridae | Troglophilinae | Troglophilus | neglectus | 1.56 | 1729811 | 1729796 | omnivor |
| Tropidischia xanthostoma | | Orthoptera | Ensifera | Rhaphidophoroidea | Rhaphidophoridae | Tropidischiinae | Tropidischia | xanthostoma | 2.2 | 1729551 | 1729550 | omnivor |
| Comicus sp. | x | Orthoptera | Ensifera | Schizodactyloidea | Schizodactylidae | Schizodactylinae | Comicus | sp. | 4.15 | | 1729847 | carnivor |
| Schizodactylus monstrosus | x | Orthoptera | Ensifera | Schizodactyloidea | Schizodactylidae | Schizodactylinae | Schizodactylus | monstrosus | 9.24 | 5101217 | 1729856 | carnivor |
| Anabropsis aptera | | Orthoptera | Ensifera | Stenopelmatoidea | Anostostomatidae | Anabropsinae | Anabropsis | aptera | 5.03 | 1725466 | 1725457 | omnivor |
| Cratomelus armatus | | Orthoptera | Ensifera | Stenopelmatoidea | Anostostomatidae | Cratomelinae | Cratomelus | armatus | 5.26 | 1725399 | 1725397 | omnivor |
| Onosandridus lanceolata | | Orthoptera | Ensifera | Stenopelmatoidea | Anostostomatidae | Anostostomatinae | Onosandridus | larvatus | 2.28 | 1725262 | 1725262 | mixed-herbivor |
| Papuaistus biroi | | Orthoptera | Ensifera | Stenopelmatoidea | Anostostomatidae | Lutosinae | Papuaistus | biroi | 5.19 | 1725112 | 1725105 | omnivor |
| Penalva lateralis | | Orthoptera | Ensifera | Stenopelmatoidea | Anostostomatidae | Anabropsinae | Penalva | lateralis | 4.1 | 1725142 | 1725136 | omnivor |
| Transaevum laudatum | x | Orthoptera | Ensifera | Stenopelmatoidea | Anostostomatidae | | Transaevum | laudatum | 3.36 | 1725335 | 1725334 | omnivor |
| Ametroides nigrifacies | | Orthoptera | Ensifera | Stenopelmatoidea | Gryllacrididae | Gryllacridinae | Ametroides | nigrifacies | 1.59 | 1726245 | 1726236 | mixed-herbivor |
| Atychogryllacris infelix | | Orthoptera | Ensifera | Stenopelmatoidea | Gryllacrididae | Gryllacridinae | Atychogryllacris | infelix | 3.73 | 1726507 | 1726506 | mixed-herbivor |

| | | | | | | | | | | | | |
|---|---|---|---|---|---|---|---|---|---|---|---|---|
| Barombogryllacris barombica | x | Orthoptera | Ensifera | Stenopelmatoidea | Gryllacrididae | Gryllacridinae | Barombogryllacris | barombica | 3.59 | 1727010 | 1727007 | mixed-herbivor |
| Capnogryllacris fumigata | x | Orthoptera | Ensifera | Stenopelmatoidea | Gryllacrididae | Hyperbaeninae | Capnogryllacris | fumigata | 4.23 | 1725889 | 1725877 | omnivor |
| Caustogryllacris podocausta | x | Orthoptera | Ensifera | Stenopelmatoidea | Gryllacrididae | Gryllacridinae | Caustogryllacris | podocausta | 2.96 | 1727188 | 1727167 | mixed-herbivor |
| Claudiagryllacris fryeri | | Orthoptera | Ensifera | Stenopelmatoidea | Gryllacrididae | Gryllacridinae | Claudiagryllacris | fryeri | 2.22 | 10024662 | 10024662 | mixed-herbivor |
| Diaphanogryllacris laeta | | Orthoptera | Ensifera | Stenopelmatoidea | Gryllacrididae | Hyperbaeninae | Diaphanogryllacris | laeta | 4.21 | 1726032 | 1726028 | omnivor |
| Diaphanogryllacris sp. | | Orthoptera | Ensifera | Stenopelmatoidea | Gryllacrididae | Hyperbaeninae | Diaphanogryllacris | sp. | 3.05 | | 1726028 | omnivor |
| Epacra aenea | | Orthoptera | Ensifera | Stenopelmatoidea | Gryllacrididae | Hyperbaeninae | Epacra | aenea | 3.12 | 1726376 | 1726375 | omnivor |
| Gryllacris bancana | | Orthoptera | Ensifera | Stenopelmatoidea | Gryllacrididae | Gryllacridinae | Gryllacris | bancana | 5.17 | 10241666 | 10290309 | mixed-herbivor |
| Kinemania ambulans | | Orthoptera | Ensifera | Stenopelmatoidea | Gryllacrididae | Gryllacridinae | Kinemania | ambulans | 2.77 | 1726227 | 1726226 | mixed-herbivor |
| Pareremus angustus | | Orthoptera | Ensifera | Stenopelmatoidea | Gryllacrididae | Gryllacridinae | Pareremus | angustus | 2.31 | 1726514 | 1726513 | mixed-herbivor |
| Sia kuhlgatzi | x | Orthoptera | Ensifera | Stenopelmatoidea | Stenopelmatidae | Stenopelmatinae | Sia | kuhlgatzi | 7.13 | 1725045 | 1725029 | omnivor |
| Sia pinguis | x | Orthoptera | Ensifera | Stenopelmatoidea | Stenopelmatidae | Stenopelmatinae | Sia | pinguis | 5.17 | 1725047 | 1725029 | omnivor |
| Stenopelmatus longispinus | x | Orthoptera | Ensifera | Stenopelmatoidea | Stenopelmatidae | Stenopelmatinae | Stenopelmatus | longispinus | 6.85 | 1725005 | 1724993 | omnivor |
| Acanthoplus discoidalis | | Orthoptera | Ensifera | Tettigonioidea | Tettigoniidae | Hetrodinae | Acanthoplus | discoidalis | 5.02 | 1687077 | 1687076 | mixed-herbivor |
| Acanthoplus longipes | | Orthoptera | Ensifera | Tettigonioidea | Tettigoniidae | Hetrodinae | Acanthoplus | longipes | 2.45 | 1687086 | 1687076 | mixed-herbivor |
| Acanthoproctus diadematus | | Orthoptera | Ensifera | Tettigonioidea | Tettigoniidae | Hetrodinae | Acanthoproctus | diadematus | 5.79 | 1684446 | 1684445 | mixed-herbivor |
| Acridoxena hewaniana | x | Orthoptera | Ensifera | Tettigonioidea | Tettigoniidae | Mecopodinae | Acridoxena | hewaniana | 5.84 | 1689037 | 1689036 | mixed-herbivor |
| Acrometopa servillea | x | Orthoptera | Ensifera | Tettigonioidea | Tettigoniidae | Phaneropterinae | Acrometopa | servillea | 2.46 | 1685590 | 1685586 | mixed-herbivor |
| Afromecopoda austera | | Orthoptera | Ensifera | Tettigonioidea | Tettigoniidae | Mecopodinae | Afromecopoda | austera | 3.63 | 1693636 | 1693631 | mixed-herbivor |
| Amyttosa mutillata | x | Orthoptera | Ensifera | Tettigonioidea | Tettigoniidae | Meconematinae | Amyttosa | mutillata | 1.26 | 1691732 | 1691728 | carnivor |
| Ancistrura nigrovittata | x | Orthoptera | Ensifera | Tettigonioidea | Tettigoniidae | Phaneropterinae | Ancistrura | nigrovittata | 2.29 | 1692724 | 1692723 | mixed-herbivor |
| Antaxius difformis | x | Orthoptera | Ensifera | Tettigonioidea | Tettigoniidae | Tettigoniinae | Antaxius | difformis | 2.49 | 1683914 | 1683907 | omnivor |
| Barbitistes fischeri | | Orthoptera | Ensifera | Tettigonioidea | Tettigoniidae | Phaneropterinae | Barbitistes | fischeri | 2.44 | 1695061 | 1695043 | forbivor |
| Barbitistes serricauda | x | Orthoptera | Ensifera | Tettigonioidea | Tettigoniidae | Phaneropterinae | Barbitistes | serricauda | 2.06 | 1695052 | 1695043 | mixed-herbivor |
| Bradyporus oniscus | | Orthoptera | Ensifera | Tettigonioidea | Tettigoniidae | Bradyporinae | Bradyporus | oniscus | 7.23 | 6544023 | 1691202 | omnivor |
| Caedicia chyzeri | x | Orthoptera | Ensifera | Tettigonioidea | Tettigoniidae | Phaneropterinae | Caedicia | chyzeri | 2.12 | 1687452 | 1687431 | mixed-herbivor |
| Championica echinus | x | Orthoptera | Ensifera | Tettigonioidea | Tettigoniidae | Pseudophyllinae | Championica | echinus | 2.47 | 1694015 | 1694014 | mixed-herbivor |
| Choeroparnops forcipatus | x | Orthoptera | Ensifera | Tettigonioidea | Tettigoniidae | Pseudophyllinae | Choeroparnops | forcipatus | 5.12 | 1686026 | 1686010 | mixed-herbivor |
| Climacoptera parallela | | Orthoptera | Ensifera | Tettigonioidea | Tettigoniidae | Pseudophyllinae | Climacoptera | parallela | 4.53 | 5096830 | 1688660 | mixed-herbivor |
| Clonia caudata | | Orthoptera | Ensifera | Tettigonioidea | Tettigoniidae | Saginae | Clonia | caudata | 4.06 | 5097435 | 1692549 | carnivor |
| Cnemidophyllum citrifolium | x | Orthoptera | Ensifera | Tettigonioidea | Tettigoniidae | Phaneropterinae | Cnemidophyllum | citrifolium | 4.43 | 1689741 | 1689739 | mixed-herbivor |
| Colossopus grandidieri | x | Orthoptera | Ensifera | Tettigonioidea | Tettigoniidae | Conocephalinae | Colossopus | grandidieri | 4.77 | 1687170 | 1687169 | omnivor |
| Conocephaloides hawaiiensis | x | Orthoptera | Ensifera | Tettigonioidea | Tettigoniidae | Conocephalinae | Conocephaloides | hawaiiensis | 3.93 | 1688372 | 1688371 | omnivor |
| Conocephalus fuscus | x | Orthoptera | Ensifera | Tettigonioidea | Tettigoniidae | Conocephalinae | Conocephalus | fuscus | 1.71 | 1683067 | 1682933 | omnivor |
| Conocephalus sp. | x | Orthoptera | Ensifera | Tettigonioidea | Tettigoniidae | Conocephalinae | Conocephalus | sp. | 1.54 | | 9397863 | omnivor |
| Corycoides abruptus | | Orthoptera | Ensifera | Tettigonioidea | Tettigoniidae | Mecopodinae | Corycoides | abruptus | 4.19 | 1694450 | 1694433 | omnivor |
| Cosmoderus erinaceus | | Orthoptera | Ensifera | Tettigonioidea | Tettigoniidae | Hetrodinae | Cosmoderus | erinaceus | 5.02 | 1687660 | 1687657 | mixed-herbivor |
| Ctenodecticus bolivari | | Orthoptera | Ensifera | Tettigonioidea | Tettigoniidae | Tettigoniinae | Ctenodecticus | bolivari | 1.8 | 1692204 | 1692203 | omnivor |
| Ctenodecticus pupulus | | Orthoptera | Ensifera | Tettigonioidea | Tettigoniidae | Tettigoniinae | Ctenodecticus | pupulus | 1.36 | 1692214 | 1692203 | omnivor |
| Cycloptera speculata | | Orthoptera | Ensifera | Tettigonioidea | Tettigoniidae | Pterochrozinae | Cycloptera | speculata | 2.46 | 5096846 | 1688722 | unknown |
| Decolya roseopicta | | Orthoptera | Ensifera | Tettigonioidea | Tettigoniidae | Meconematinae | Decolya | roseopicta | 1.34 | 1683650 | 1683645 | carnivor |
| Decticus verrucivorus | x | Orthoptera | Ensifera | Tettigonioidea | Tettigoniidae | Tettigoniinae | Decticus | verrucivorus | 4.05 | 1690685 | 1690683 | omnivor |

| | | | | | | | | | | | | |
|---|---|---|---|---|---|---|---|---|---|---|---|---|
| Ephippiger provincialis | x | Orthoptera | Ensifera | Tettigonioidea | Tettigoniidae | Bradyporinae | Ephippiger | provincialis | 4.69 | 1683971 | 1683970 | omnivor |
| Eubliastes adustus | | Orthoptera | Ensifera | Tettigonioidea | Tettigoniidae | Pseudophyllinae | Eubliastes | adustus | 3.74 | 1694458 | 1694452 | mixed-herbivor |
| Euconchophora spinigera | x | Orthoptera | Ensifera | Tettigonioidea | Tettigoniidae | Conocephalinae | Euconchophora | spinigera | 2.56 | 1686547 | 1686546 | omnivor |
| Eupholidoptera smyrnensis | | Orthoptera | Ensifera | Tettigonioidea | Tettigoniidae | Tettigoniinae | Eupholidoptera | smyrnensis | 3.6 | 1690278 | 1690233 | omnivor |
| Gampsocleis glabra | x | Orthoptera | Ensifera | Tettigonioidea | Tettigoniidae | Tettigoniinae | Gampsocleis | glabra | 3.12 | 1694547 | 1694531 | omnivor |
| Geonotus vittatus | | Orthoptera | Ensifera | Tettigonioidea | Tettigoniidae | Pseudophyllinae | Geonotus | vittatus | 2.7 | 1692504 | 1692503 | mixed-herbivor |
| Gonamytta occidentalis | x | Orthoptera | Ensifera | Tettigonioidea | Tettigoniidae | Meconematinae | Gonamytta | occidentalis | 1.26 | 1690121 | 1690117 | carnivor |
| Gymnoproctus abortivus | | Orthoptera | Ensifera | Tettigonioidea | Tettigoniidae | Hetrodinae | Gymnoproctus | abortivus | 4.74 | 1685144 | 1685139 | mixed-herbivor |
| Hemisaga denticulata | | Orthoptera | Ensifera | Tettigonioidea | Tettigoniidae | Austrosaginae | Hemisaga | denticulata | 4.87 | 1693189 | 1693180 | carnivor |
| Hetrodes pupus | | Orthoptera | Ensifera | Tettigonioidea | Tettigoniidae | Hetrodinae | Hetrodes | pupus | 5.56 | 1690937 | 1690936 | mixed-herbivor |
| Hexacentrus mundus | x | Orthoptera | Ensifera | Tettigonioidea | Tettigoniidae | Hexacentrinae | Hexacentrus | mundus | 2.76 | 1688473 | 1688450 | carnivor |
| Incertana decorata | x | Orthoptera | Ensifera | Tettigonioidea | Tettigoniidae | Tettigoniinae | Incertana | decorata | 2.5 | 7562760 | 1683749 | omnivor |
| Insara elegans | | Orthoptera | Ensifera | Tettigonioidea | Tettigoniidae | Phaneropterinae | Insara | elegans | 1.59 | 1694608 | 1694582 | mixed-herbivor |
| Jamaicana subguttata | x | Orthoptera | Ensifera | Tettigonioidea | Tettigoniidae | Pseudophyllinae | Jamaicana | subguttata | 3.74 | 1683510 | 1683508 | mixed-herbivor |
| Leproscirtus granulosus | x | Orthoptera | Ensifera | Tettigonioidea | Tettigoniidae | Mecopodinae | Leproscirtus | granulosus | 2.26 | 1686644 | 1686641 | mixed-herbivor |
| Lesina ensifera | x | Orthoptera | Ensifera | Tettigonioidea | Tettigoniidae | Conocephalinae | Lesina | ensifera | 9.29 | 1687273 | 1687266 | omnivor |
| Macroxiphus sumatranus | x | Orthoptera | Ensifera | Tettigonioidea | Tettigoniidae | Conocephalinae | Macroxiphus | sumatranus | 4.22 | 1683374 | 1683373 | omnivor |
| Meconema meridionale | x | Orthoptera | Ensifera | Tettigonioidea | Tettigoniidae | Meconematinae | Meconema | meridionale | 0.99 | 1690429 | 1690427 | carnivor |
| Metaplastes ornatus | x | Orthoptera | Ensifera | Tettigonioidea | Tettigoniidae | Phaneropterinae | Metaplastes | ornatus | 2.07 | 1687733 | 1687731 | mixed-herbivor |
| Metrioptera saussuriana | x | Orthoptera | Ensifera | Tettigonioidea | Tettigoniidae | Tettigoniinae | Metrioptera | saussuriana | 2.44 | 1685554 | 1685533 | omnivor |
| Moncheca pretiosa | x | Orthoptera | Ensifera | Tettigonioidea | Tettigoniidae | Conocephalinae | Moncheca | pretiosa | 4.18 | 1687790 | 1687780 | omnivor |
| Mortoniellus karnyi | x | Orthoptera | Ensifera | Tettigonioidea | Tettigoniidae | Lipotactinae | Mortoniellus | karnyi | 4.46 | 1683609 | 1683603 | carnivor |
| Neobarrettia sinaloae | | Orthoptera | Ensifera | Tettigonioidea | Tettigoniidae | Listroscelidinae | Neobarrettia | sinaloae | 4.43 | 1687919 | 1687903 | carnivor |
| Neocallicrania bolivarii | x | Orthoptera | Ensifera | Tettigonioidea | Tettigoniidae | Bradyporinae | Neocallicrania | bolivarii | 3.93 | 4403242 | 1683452 | omnivor |
| Odontolakis sexpunctata | | Orthoptera | Ensifera | Tettigonioidea | Tettigoniidae | Conocephalinae | Odontolakis | sexpunctata | 7.08 | 1683399 | 1683397 | omnivor |
| Odontura aspericauda | x | Orthoptera | Ensifera | Tettigonioidea | Tettigoniidae | Phaneropterinae | Odontura | aspericauda | 1.82 | 5096140 | 1683719 | mixed-herbivor |
| Paracaedicia verrucosa | | Orthoptera | Ensifera | Tettigonioidea | Tettigoniidae | Phaneropterinae | Paracaedicia | verrucosa | 2.29 | 1686707 | 1686695 | mixed-herbivor |
| Phisis holdhausi | | Orthoptera | Ensifera | Tettigonioidea | Tettigoniidae | Meconematinae | Phisis | holdhausi | 1.5 | 1686844 | 1686834 | carnivor |
| Phlugidia usambarica | | Orthoptera | Ensifera | Tettigonioidea | Tettigoniidae | Meconematinae | Phlugidia | usambarica | 1.01 | 1685458 | 1685457 | carnivor |
| Phlugis teres | | Orthoptera | Ensifera | Tettigonioidea | Tettigoniidae | Meconematinae | Phlugis | teres | 0.95 | 1694298 | 1694262 | carnivor |
| Phyllophora cheesmanae | x | Orthoptera | Ensifera | Tettigonioidea | Tettigoniidae | Phyllophorinae | Phyllophora | cheesmanae | 3.3 | 5097087 | 1690322 | mixed-herbivor |
| Platystolus martinezii | | Orthoptera | Ensifera | Tettigonioidea | Tettigoniidae | Bradyporinae | Platystolus | martinezii | 4.78 | 5097516 | 8154847 | omnivor |
| Poecilimon jonicus | x | Orthoptera | Ensifera | Tettigonioidea | Tettigoniidae | Phaneropterinae | Poecilimon | jonicus | 1.6 | 1688178 | 1688084 | other |
| Prohimerta fujianensis | x | Orthoptera | Ensifera | Tettigonioidea | Tettigoniidae | Phaneropterinae | Prohimerta | fujianensis | 1.56 | 1686594 | 1686587 | mixed-herbivor |
| Pseudophyllanax imperialis | | Orthoptera | Ensifera | Tettigonioidea | Tettigoniidae | Mecopodinae | Pseudophyllanax | imperialis | 2.81 | 1692321 | 1692320 | mixed-herbivor |
| Pycnogaster graellsii | | Orthoptera | Ensifera | Tettigonioidea | Tettigoniidae | Bradyporinae | Pycnogaster | graellsii | 4.36 | 1694873 | 1694869 | omnivor |
| Ruspolia nitidula | x | Orthoptera | Ensifera | Tettigonioidea | Tettigoniidae | Conocephalinae | Ruspolia | nitidula | 3.33 | 5096614 | 1686351 | omnivor |
| Sanaa intermedia | x | Orthoptera | Ensifera | Tettigonioidea | Tettigoniidae | Pseudophyllinae | Sanaa | intermedia | 5.32 | 1690627 | 1690621 | mixed-herbivor |
| Sasima aequalis | | Orthoptera | Ensifera | Tettigonioidea | Tettigoniidae | Phyllophorinae | Sasima | aequalis | 4.28 | 1684888 | 1684881 | mixed-herbivor |
| Sathrophyllia femorata | | Orthoptera | Ensifera | Tettigonioidea | Tettigoniidae | Pseudophyllinae | Sathrophyllia | femorata | 3.71 | 1693220 | 1693217 | mixed-herbivor |
| Sureyaella bella | x | Orthoptera | Ensifera | Tettigonioidea | Tettigoniidae | Tettigoniinae | Sureyaella | bella | 1.72 | 1689707 | 1689706 | omnivor |
| Triencentrus atrosignatus | | Orthoptera | Ensifera | Tettigonioidea | Tettigoniidae | Pseudophyllinae | Triencentrus | atrosignatus | 3.59 | 1689222 | 1689216 | mixed-herbivor |

| | | | | | | | | | | | | |
|---|---|---|---|---|---|---|---|---|---|---|---|---|
| Tympanophyllum atroterminatum | x | Orthoptera | Ensifera | Tettigonioidea | Tettigoniidae | Pseudophyllinae | Tympanophyllum | atroterminatum | 2.96 | 1684679 | 1684653 | mixed-herbivor |
| Typophyllum trapeziforme | | Orthoptera | Ensifera | Tettigonioidea | Tettigoniidae | Pterochrozinae | Typophyllum | trapeziforme | 2.1 | 1693592 | 1693561 | unknown |
| Xiphidiola dispersa | x | Orthoptera | Ensifera | Tettigonioidea | Tettigoniidae | Meconematinae | Xiphidiola | dispersa | 1 | 1692085 | 1692067 | carnivor |
| Zaprochilus australis | | Orthoptera | Ensifera | Tettigonioidea | Tettigoniidae | Zaprochilinae | Zaprochilus | australis | 1.61 | 1687653 | 1687649 | other |
| Dinocras cephalotes | x | Outgroup | Outgroup | Outgroup | Outgroup | Outgroup | Dinocras | cephalotes | 0.59 | 2005247 | 2005240 | omnivor |
| Forficula aetolica | x | Outgroup | Outgroup | Outgroup | Outgroup | Outgroup | Forficula | aetolica | 0.19 | 4393040 | 1419844 | carnivor |
| Grylloblatta bifratilectra | x | Outgroup | Outgroup | Outgroup | Outgroup | Outgroup | Grylloblatta | bifratilectra | 1.22 | 1407051 | 1407042 | omnivor |
| Periplaneta americana | x | Outgroup | Outgroup | Outgroup | Outgroup | Outgroup | Periplaneta | americana | 2.31 | 2000152 | 1993899 | omnivor |
| Xylica oedematosa | x | Outgroup | Outgroup | Outgroup | Outgroup | Outgroup | Xylica | oedematosa | 2.79 | 1415852 | 1415847 | forbivor |
| Zorotypus caudelli | x | Outgroup | Outgroup | Outgroup | Outgroup | Outgroup | Zorotypus | caudelli | 0.15 | 1407016 | 1406991 | forbivor |
